# Enviromic assembly increases accuracy and reduces costs of the genomic prediction for yield plasticity

**DOI:** 10.1101/2021.06.04.447091

**Authors:** Germano Costa-Neto, Jose Crossa, Roberto Fritsche-Neto

## Abstract

Quantitative genetics states that phenotypic variation is a consequence of genetic and environmental factors and their subsequent interaction. Here, we present an enviromic assembly approach, which includes the use of ecophysiology knowledge in shaping environmental relatedness into whole-genome predictions (GP) for plant breeding (referred to as E-GP). We propose that the quality of an environment is defined by the core of environmental typologies (envirotype) and their frequencies, which describe different zones of plant adaptation. From that, we derive markers of environmental similarity cost-effectively. Combined with the traditional genomic sources (e.g., additive and dominance effects), this approach may better represent the putative phenotypic variation across diverse growing conditions (i.e., phenotypic plasticity). Additionally, we couple a genetic algorithm scheme to design optimized multi-environment field trials (MET), combining enviromic assembly and genomic kinships to provide in-silico realizations of the future genotype-environment combinations that must be phenotyped in the field. As a proof-of-concept, we highlight E-GP applications: (1) managing the lack of phenotypic information in training accurate GP models across diverse environments and (2) guiding an early screening for yield plasticity using optimized phenotyping efforts. Our approach was tested using two non-conventional cross-validation schemes to better visualize the benefits of enviromic assembly in sparse experimental networks. Results on tropical maize show that E-GP outperforms benchmark GP in all scenarios and cases tested. We show that for training accurate GP models, the genotype-environment combinations’ representativeness is more critical than the MET size. Furthermore, we discuss theoretical backgrounds underlying how the intrinsic envirotype-phenotype covariances within the phenotypic records of (MET) can impact the accuracy of GP and limits the potentialities of predictive breeding approaches. The E-GP is an efficient approach to better use environmental databases to deliver climate-smart solutions, reduce field costs, and anticipate future scenarios.

## 1 INTRODUCTION

Environmental changing scenarios challenge agricultural research to deliver climate-smart solutions in a time-reduced and cost-effective manner (Tigchelaar *et al*., 2018; Ramírez-Villegas *et al* 2020; Cortés *et al*., 2020). Characterizing crop growth conditions is crucial for this purpose (Xu, 2016), allowing a deeper understanding of how the environment shapes past, present, and future phenotypic variations (e.g., Ramírez-Villegas *et al*. 2018; Heinemann *et al*., 2019; Cooper *et al*., 2014; de los Campos *et al*., 2020; Costa-Neto *et al*., 2021b; Antolin et al., 2021). For plant breeding research, mostly based on selecting the best-evaluated genotypes for a target population of environments (TPE), this approach is useful to discriminate genomic and non-genomic sources of crop adaptation. Thus, the concept of ‘envirotyping’ (environmental + typing, Cooper *et al*., 2014; Xu, 2016) emerges to establish the quality of a given environment in the delivery of quality phenotypic records, mostly to train accurate predictive breeding approaches capable of guiding the selection of most productive and adapted genotypes (Resende *et al*., 2020; Costa Neto *et al*., 2021a; Crossa *et al*., 2021).

From envirotyping, it is possible to check the quality of a certain environment, which is directly related to how the observed growing conditions in a particular field trial could be related to the most frequent environment-types (envirotypes) that occur in the breeding program TPE or target region (e.g., Heinemann et al., 2019; Cooper *et al*., 2021; Antolin *et al*., 2021). In agricultural research, the quality of a certain environment is directly related to how it can limit the expression of the genetic potential of the certain crop for a certain trait, such as suggested by the movement called ‘School of de Wit’ since 1965 (see Bouman *et al*., 1996). Thus, for the plant breeding research, this is also direct factors such as genotype × environment interaction (e.g., Allard, 1964; Finlay and Wilkinson, 1963) and its implications of how the target germplasm under selection (or testing) can perform across the target growing conditions in which the candidate cultivars will be cropped.

Prediction-based tools have leveraged modern plant breeding research to an extent in which phenotyping is still required (Crossa *et al*., 2017), although prediction-based tools and simulations can support more comprehensive and faster selection decisions (Galli *et al*., 2020; Cooper *et al*., 2021; Crossa *et al*., 2021). One of the most widely used predictive tools is the whole-genome prediction (GP, Meuwissen *et al*., 2001), developed and validated for several crop species and application scenarios (Crossa *et al*., 2017; Voss-Fels *et al*., 2019), such as the selection among populations and the prediction of the performance of single-crosses across multiple environments. For the latter, the most important use of GP mostly relies on the better use of the available phenotyping records and large-scale easy-managed genomic information to expand the spectrum of evaluated single-crosses in silico (Messina *et al*., 2018; Rogers *et al*., 2021). Those phenotypic records (e.g., grain yield and plant height) are collected from existing field trials that experience a diverse set of growing conditions, carrying within them an intrinsic environment-phenotype covariance. Consequently, the GP has a limited accuracy under multiple-environment testing (MET) due to genotype × environment interaction (G×E) (Crossa *et al*., 2017), meaning that each genotype has a differential response for each environmental factor that assembles what we call ‘environment’ (time interval across crop lifetime involving a specific geographic location and agronomic practice for a particular crop). Therefore, novel ways to include environmental data (Heslot *et al*., 2014; Jarquín *et al*., 2014; Ly *et al*., 2018; Millet *et al*., 2019; Gillberg *et al*., 2019; Costa-Neto *et al*., 2021a) and process-based crop growth models (CGM) (Messina *et al*., 2018; Toda *et al*. 2020; Robert *et al*., 2020; Cooper *et al*., 2021) in GP are considered the best pathways to fix it. Most of the success achieved by such approaches lies in a better understanding of the visible ecophysiology interplay between genomics and environment variation (Gage *et al*., 2017; Li *et al*., 2018; Guo *et al*., 2020; Costa-Neto *et al*., 2021b).

The explicit integration of enviromic and genomic sources is an easy way to lead GP to a wide range of novel applications (Crossa *et al*., 2021), such as improving the predictive ability for untested growing conditions (Guo *et al*., 2020; de los Campos *et al*., 2020; Jarquín *et al*., 2020; Costa-Neto *et al*., 2021a), to optimize MET networks and to screen genotype-specific reaction-norms (Ly *et al*., 2018; Millet *et al*., 2019). This is excellent progress for predictive breeding (i.e., the range of prediction-based selection tools for crop improvement) and accelerating research pipelines to deliver higher yields and adapted genotypes for target scenarios. However, most of the current studies on this topic vary in accuracy and applicability, mostly due to (1) the processing protocols used to translate the raw-data into explicit environmental covariables (ECs) with biological meaning in explaining G×E over complex traits, (2) the lack of a widely-used envirotyping pipeline that, not only supports the design of field trials, but also increases the accuracy of the trained GP models and, in addition, (3) for CGM, a possible limitation is the increased demand for the phenotyping of additional intermediate phenotypes (i.e., biomass accumulation and partitioning, specific leaf area), which can involve managed iso-environments and expert knowledge in crop modeling (Cooper *et al*., 2016; Toda *et al*., 2020; Robert *et al*., 2020). The latter can be expensive or difficult for plant research programs in developing countries, which generally have low budgets to increase the phenotyping network and install environmental sensors. In addition, most developing countries are located in regions where environments are subject to a broader range of stress factors (e.g., heat stress).

Therefore, here we revisit Shelford’s Law (Shelford, 1931) and other ecophysiology concepts that can provide the foundations for translating raw-environmental information into an enviromic source for predictive breeding, hereafter denominated as *enviromic assembly*. The benefits of using the so-called ‘enviromics-aided GBLUP’ (E-GP) under existing experimental networks are presented, followed by the E-GP application to optimize field-based phenotyping. Finally, we benchmark E-GP with the traditional genomic-best unbiased prediction (GBLUP) to discuss the benefits of enviromic data to reproduce G×E patterns and provide a virtual screening for yield plasticity.

## 2 MATERIAL AND METHODS

The material methods are organized in the following manner: First, we briefly address the concepts underlying the novel approach of *enviromic assembly* inspired by Shelford’s Law. The data sets are then presented, along with the statistical models and prediction scenarios used to show the benefits of large-scale environmental information in GP across multi-environment trials (MET). Finally, we present a scheme to optimize phenotyping efforts in training GP over MET and support the screening for maize single-crosses’ yield plasticity.

### 2.1 Theory: adapting the Shelford Law of Minimum

Consider two experimental networks (MET) of the same target population of environments (TPE, e.g., the different locations, years, and crop management) under different environmental gradients due to year or location variations (Fig.1). For two genotypes evaluated under both conditions (G1, G2), the potential genetic-specific phenotypic plasticity (Allard and Bradshaw, 1964) (curves) is expressed as different reaction-norms (dotted lines), resulting in distinct observable G×E patterns (Fig.1a-b). In the former MET (Fig.1a), both genotypes experience a wider range of possible growing conditions (large interval between the two vertical solid lines), which result in an intricate G×E pattern (crossover). Conversely, in the latter MET (Fig.1b), the same genotypes experience a reduced range of growing conditions yet lead to a simple G×E pattern (non-crossover). It is feasible to conclude that, although the genetic variation is essential for modeling potential phenotypic plasticity of genotypes (curves, Fig.1a-c), the diversity of environmental growing conditions dictates the observable G×E patterns (Bradshaw, 1965). Thus, the GP platforms for MET may be unbiased with no diversity, and the quality of environments is not considered.

**Fig. 1.**
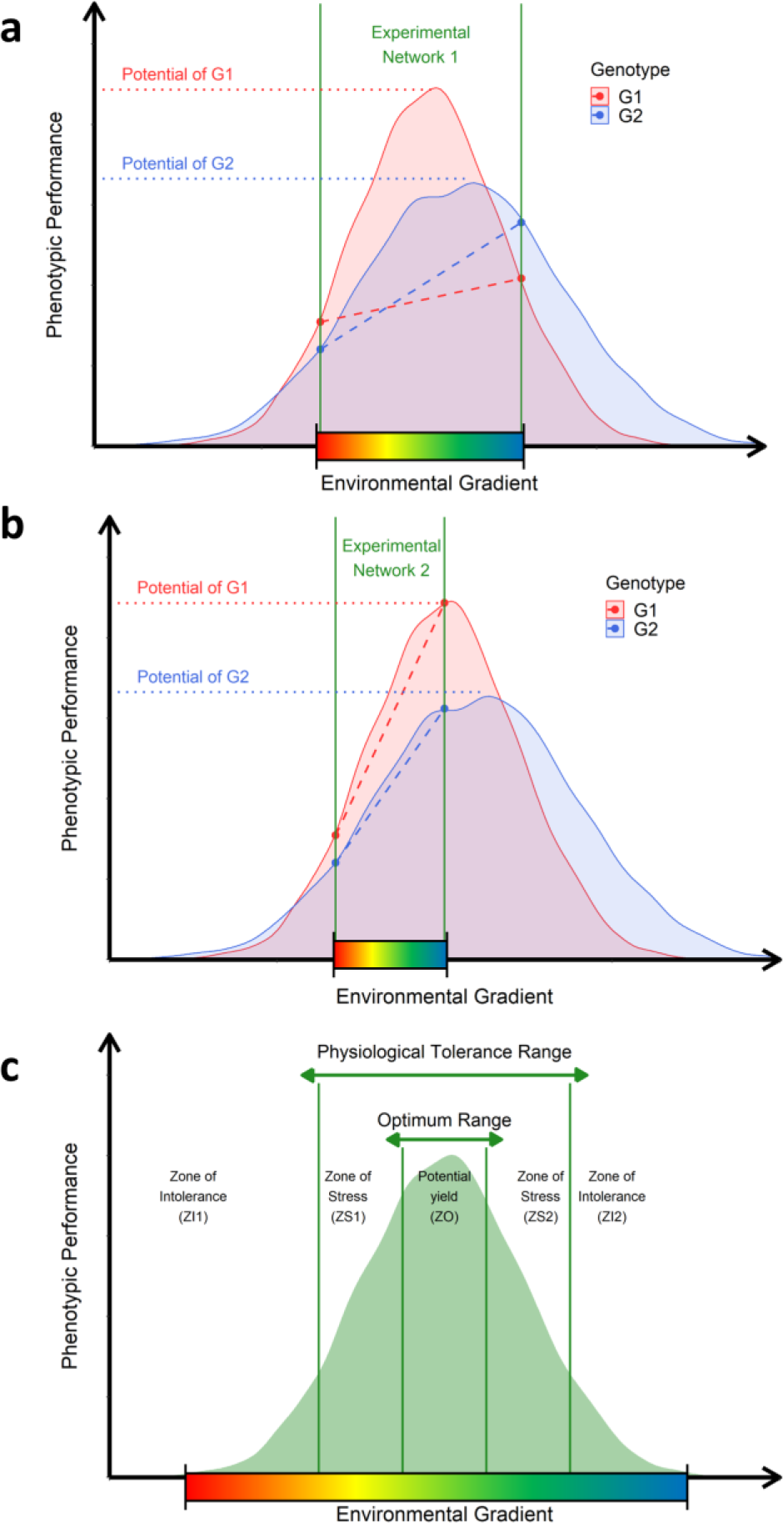
Ecophysiological insights to translate raw-environmental data into enviromic sources. **a**. Representation of an experimental network involving an unknown number of environments from a theoretical TPE and two genotypes (G1 and G2). The range of the environmental gradient is delimited by the space between the two vertical green lines. Each genotype has a nonlinear function describing the genetic limits of their phenotypic plasticity (curves) and genetic potential (horizontal dotted lines) of a given trait. Diagonal dotted lines denote the observed reaction-norm experienced by those genotypes; **b**. representation of a second experimental network involving the same genotypes, but different environments were sampled from the theoretical TPE. **c**. adaptation of Shelford’s Law of Tolerance, describing the cardinal (or biological) genetic limits (vertical green lines) to determine the amount of the factor that results in different adaptation zones. Across these zones, crop performance is described by zones of stress caused by deficit or excess (physiological tolerance range) and zones of optimal growing conditions that allow the plants to express the genetic potential for a given trait (optimum range). The core of possible environmental variations contemplated as putative phenotypic plasticity for a given genotype, germplasm, or crop species.

Approaches such as CGM try to reproduce the phenotypic plasticity curves, while benchmark reaction-norm models try to reproduce the observable reaction-norm. Both approaches can achieve adequate results, although we have observed that (1) CGM demands greater phenotyping efforts to train computational approaches capable of reproducing the *achievable* phenotypic plasticity from a reduced core of phenotypic records from field trials at near-iso environments (e.g., well-watered conditions versus water-limited conditions for the same planting date and management), (2) CGM demands additional programing efforts, which, for some regions or crops, can be expensive and limit the applicability of the method, (3) adequate reaction-norm models over well-designed phenotyping platforms are not a reality for certain regions of the world with limited resources to invest in precision phenotyping efforts.

We understand that Shelford’s Law of Tolerance (Shelford, 1931) is suitable to explain how the environment drives plant plasticity and can be incorporated into the traditional GP platforms in a cost-effective way (Fig.1c). It states that a target population’s adaptation is modulated as a certain range of minimum, maximum and optimum threshold limits achieved over an environment gradient (vertical solid green lines). The genotypes’ potential phenotypic plasticity (curves) is not regarded as a linearized reaction-norm variation across an environmental gradient (Arnold *et al*., 2019). Instead, it is the distribution of possible phenotypic expressions dictated by the cardinal thresholds for each biophysical factor with ecophysiological relevance. Therefore, crops may experience stressful conditions due to the excess or lack of a target environmental factor, depending on the cardinal thresholds (vertical solid green lines in Fig.1c), which also rely on some key development stages germplasm-specific characteristics (e.g., tropical maize versus temperate maize). Consequently, the expected variation of environmental conditions across different field trials results from a series of environment-types (envirotypes) acting consistently yet varying in impact depending on the genetic-specific sensibility. The quality of a certain growing condition depends on the balance between crop necessity and resource availability, which involves *constant effects*, such as the type of treatments in a trial (e.g., fertilizer inputs) and *transitory effects* variables, such as weather events (e.g., heat-stress).

From these concepts, we observe that with the use of envirotyping (e.g., typing the profiles of a particular environment), the environment part of the G×E pattern can be visualized based on the shared frequency of envirotypes among different field trials. Thus, the enviromic of a certain experimental network or TPE (the core of possible growing conditions) can be mathematically assembled by (1) collecting large-scale environmental data, (2) processing this raw data in envirotyping entries for each real or virtual environment, and (3) processing these envirotyping-derived entries to achieve theoretical relatedness between the buildup of different environments from the shared frequency of envirotypes. Thus, the expected envirotypes can be designed relying on the adaptation zones inspired by the model proposed here, based on Shelford’s Law, in which we can envisage the process of deriving environmental covariables for GP into an ecophysiological-smart way.

### 2.2 Proof-of-concept data sets

This study used maize as a proof-of-concept crop due to its importance for food security in developed and developing regions. Two data sets of maize hybrids (single-crosses of inbreed lines) from different germplasm sources developed under tropical conditions in Brazil (hereafter referred to as Multi-Regional and N-level) were used. Both data sets involve phenotypic records of grain yield (Mg per ha) collected across multiple environments. Details on the experimental design, cultivation practices, and fundamental statistical analysis are given in Bandeira e Souza *et al*. (2017) and Alves *et al*. (2019). Below we provide a short description of the number of genotypes and environments tested and the nature of this study’s genotyping data.

#### 2.2.1 Multi-Regional Set

The so-called “Multi-Regional set” is based on the germplasm developed by the Helix Seeds Company (HEL) in South America. It includes 247 maize lines evaluated in 2015 in five locations in three regions of Brazil (Supplementary Table 1). Genotypes were obtained using the Affymetrix Axiom Maize Genotyping Array containing 616 K SNPs (single-nucleotide polymorphisms) (Unterseer *et al*., 2014). Only SNPs with a minor allele frequency > 0.05 were considered. Finally, a total of 52,811 high-quality SNPs that achieved the quality control level were used in further analysis.

#### 2.2.2 N-level set

The so-called “N-level set” is based on the germplasm developed by the Luiz de Queiroz College of Agriculture of the University of São Paulo (USP), Brazil. A total of 570 tropical maize hybrids were evaluated across eight environments, involving an arrangement of two locations, two years, and two nitrogen levels (Supplementary Table 2). This study’s sites involved two distinct edaphoclimatic patterns, i.e., Piracicaba (Atlantic forest, clay soil) and Anhumas (savannah, silt–sandy soil). In each site, two contrasting nitrogen (N) fertilization levels were managed. One experiment was conducted under ideal N conditions and received 30 kg ha^−1^ at sowing, along with 70 kg ha^−1^ in a coverage application at the V8 plant stage. That is the main recommendation for fertilization in tropical maize growing environments in Brazil. In contrast, the second experiment under low N conditions received only 30 kg ha^−1^ of N at sowing, resulting in an N-limited growing condition. This set’s genotypes were also obtained using the Affymetrix Axiom Maize Genotyping Array containing 616 K SNPs (Unterseer *et al*., 2014) and minor allele frequency > 0.05. At the end of this process, a total of 54,113 SNPs were considered in the GP modeling step.

### 2.3 Envirotyping Pipeline

Below, we present the methods used for data collection, data processing, and implementing what we call ‘enviromic assembly’. This envirotyping pipeline was developed using the functions of the R package *EnvRtype* (Costa-Neto *et al*., 2021) and is available at no cost. All codes for running the next steps are given in https://github.com/gcostaneto/EGP.

#### 2.3.1 Environmental sensing (data collection)

In this study, environmental information was used for the main abiotic plant-environment interactions related to daily weather, soil type, and crop management (available only for N-level set). Daily weather information was collected from NASA POWER (Sparks, 2018) and consisted of eight variables: rainfall (P, mm day^-1^), maximum air temperature (TMAX, °C day^-1^), minimum air temperature (TMIN, °C day^-1^), average air temperature (TAVG, °C day^-1^), dew point temperature (TDEW, °C day^-1^), global solar radiation (SRAD, MJ per m²), wind speed at 2 meters (WS, m s^-1^ day^-1^) and relative air humidity (RH, % day^-1^). Elevation above sea level was obtained from NASA’s Shuttle Radar Topography Mission (SRTM). Both sources were imported into R statistical-computational environments using the functions and libraries organized within the *EnvRtype* package (Costa-Neto *et al*., 2021b). A third GIS database was used to import soil types from Brazilian soil classification provided by EMBRAPA and available at https://github.com/gcostaneto/EGP.

#### 2.3.2 Data Processing

Quality control was adopted by removing variables outside the mean ± three standard deviation and repeated columns. After checking for outliers, the daily weather variables were used to model ecophysiological interactions related to soil-plant-atmosphere dynamics. The thermal-radiation interactions computed potential atmospheric evapotranspiration (ET0) following the Priestley-Taylor method. The slope of the saturation vapor pressure curve (SPV) and vapor pressure deficit (VPD) was computed as given in the FAO manual (Allen *et al*., 1998). An FAO-based generic function was used to estimate crop development as a function of days after emergence (DAE). We assume a 3-segment leaf growing function to estimate the crop canopy coefficient (Kc) of evapotranspiration using the following Kc values: Kc_1_ (0.3), Kc_2_ (1.2), Kc_3_ (0.35), equivalent to the water demand of tropical maize for initial phases, reproduction phases, and end-season stages, respectively. Using the same 3-segment function, we estimate the crop canopy using a leaf area index (LAI) of LAI = 0.7 (initial vegetative phases), LAI = 3.0 (maximum LAI for tropical maize growing conditions observed in our fields), and LAI = 2.0 (LAI tasseling stage). We computed the daily crop evapotranspiration (ETc) estimated by the product between ET0 and the Kc from those two estimations. Then, we computed the difference between daily precipitation and crop evapotranspiration as P-ETc.

The apparent photosynthetic radiation intercepted by the canopy (aPAR) was computed following aPAR=SRAD×(1-exp(-k×LAI)), where k is the coefficient of canopy, considered as 0.5. Water deficiency was computed using the atmospheric water balance between input (precipitation) and output of atmospheric demands (crop evapotranspiration). The effect of temperature on the radiation use efficiency (F_RUE_) was described by a three-segment function based on cardinal temperatures for maize, using the cardinal temperatures 8°C (Tb_1_, base lower), 30°C (To_1_, base optimum), 37°C (To_2_, upper optimum) and 45°C (Tb_2_, base upper). This function assumes values from 0 to 1, depending on: F_RUE_= 0 if T_AVG_ ≤ Tb_1_; F_RUE_ = (T_AVG_ -Tb_1_)/(To_1_-Tb_1_) if Tb_1_ < T_AVG_ < To_1_; F_RUE_ = 1 if To_1_ < T_AVG_ < To_2_; F_RUE_ = (Tb_2_ – T_AVG_)/(Tb_2_ – To_2_) if To_2_ < T_AVG_ < Tb_2_; and F_RUE_ = 0 if T_AVG_ > Tb_2_.

Finally, we sampled each piece of weather and ecophysiological information across five-time intervals in the crop lifetime: from emergence to the appearance of the first leaf (V1, 14 DAE), from V1 to the fourth leaf (V4, 35 DAE), from V4 to the tasseling stage (VT, 65 DAE), from VT to the kernel milk stage (R3, 90 DAE) and from R3 to physiological maturity (120 DAE), in which DAE stands for days after emergence.

#### 2.3.3 Enviromic assembly using typologies (T matrix)

The raw envirotyping data were used to assemble markers for environmental similarity, depending on the group of the ECs. The first group of ECs involves the transitory effect variables, which vary in the frequency of occurrence, depending on the crop development cycle. Thus, we design the expected envirotypes using the number of inputs required to lead crops in at least three levels of adaptation: (1) stress by deficit, (2) optimum growing conditions, and (3) stress by excess. These levels were defined using cardinal thresholds or frequency tables concerning the growing conditions archived in the experimental network range. Then, from having reviewed the literature, we consider the intervals for thermal-related variables: 0°C to 9°C (death), 9.1°C to 23°C (stress by deficit), 23.1°C to 32°C (optimum growing conditions), 32.1°C to 45°C (stress by excess) and 45°C to ∞°C (death). We computed the classes for accumulated prediction according to our agronomic expertise on rainfall requirements for tropical maize growing environments: 0mm to 10mm, 10.1mm to 20mm, and 20.1mm to ∞ mm. For crop evapotranspiration (ETc), we assume the envirotypes 0-6 mm.day-1, 7-10 mm.day-1, 10-15 mm.day-1 and 16 to ∞ mm.day-1. Finally, for F_RUE_, we assume the envirotypes based on the following adaptation zones: impact from 0% to 25% (0-0.25), from 26% to 50% (0.26-0.50), 51% to 75% (0.51-0.75) and 76% to 100% (0.76-1.0). We preferred to adopt a simple discretization for the remaining variables using a histogram of percentiles (0-25%, 26-50%, 51-75%, 75-100%) of occurrence for a target envirotype.

The second group involves constant effect variables. In this group, we consider the elevation, crop management, and soil classification in each environment. Soil information was entered as an incidence matrix (0 or 1) based on each environment’s occurrence. In addition, for the N-level set, nitrogen input levels were computed as two discrete classes: ideal N = 10 and low N = 30; we entered this same incidence matrix for soil information. Because both sets have a gradient for elevation, we used a histogram of percentiles (0-25%, 26-50%, 51-75%, 75-100%) as in the transitory group of variables. Finally, each designed envirotype × time interval frequency was used as a qualitative marker of environmental relatedness (the hereafter **T** matrix, from typologies).

#### 2.3.4 Assembly of W matrix using quantile covariables (benchmark EC matrix)

The quantitative descriptors of environmental relatedness are the most common method to include environmental information in GP studies considering reaction-norms. Jarquín *et al*. (2014) proposed the creation of the so-called environmental relatedness kinship (***K***_E_) carried out with a matrix of quantitative environmental covariables (**W** matrix, thus we refer to this environment kinship as ***K***_E,W_). Here, this pattern of similarity in ***K***_E,W_ was captured using percentile values (25%; 50%, and 75%) at each of the five-time intervals of development, as suggested by Morais-Júnior *et al*. (2018) and expanded by Costa-Neto *et al*., (2021a). We found 255 and 307 quantitative descriptors for the Multi-Regional and N-level sets at the end of the process, respectively. In this study, we used this ***K***_E,W_ as a benchmark method to test the effectiveness of the ***K***_E,T_ matrix and the total absence of environmental information (baseline genomic model without environmental information, see section 2.4.1).

### 2.4 Statistical Models

From a baseline additive-dominant multi-environment GBLUP (section 2.4.1), we tested four other models, created with the inclusion of two types of enviromic assembly (**T** or **W**) and structures for G×E effects. More details about each statistical model are provided in the next subsections. All kernel models were fitted using the BGGE R package (Granato *et al*., 2018) using 15,000 iterations, with 2,000 used as burn-in and using a thinning of 10. This package was used due to the following aspects: (1) is an accurate open-source software and; (2) can accommodate many kernels in a computation-efficient way.

#### 2.4.1 Baseline additive-dominant GBLUP

The baseline model includes a fixed intercept for each environment and random genetic variations (additive and dominance). We will refer to this model as GBLUP, which was modeled as an overall main effect plus a genomic-by-environment deviation (the so-called G+GE model, Bandeira e Souza *et al*., 2017), as follows:

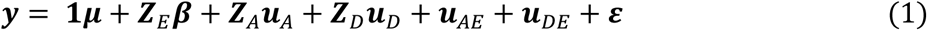

where ***y*** = [***y***_***1***_, ⋯, ***y***_***n***_]^′^ is the vector of observations collected in each of the *q* environments with hybrids and ***1μ*** + ***Z***_E_***β*** is the general mean and the fixed effect of the environments with incidence matrix ***Z***_*E*_. Genetic variations are modeled using the main additive effects (***u***_A_), with ***u***_A_ ∼ 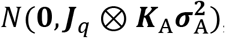, plus a random dominance variation (***u***_D_), with ***u***_D_ ∼ 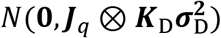, where 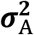 and 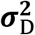 are the variance component for additive and dominance deviation effects; ***Z***_A_ and ***Z***_D_ are the incidence matrix for the same effects (absence=0, presence=1), ***J***_*q*_ is a *q*×*q* matrix of 1s and ⊗ denotes the Kronecker Product. G×E effects are modeled using a block diagonal (BD) matrix of the genomic effects, built using 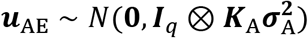 and 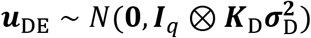, in which ***I***_*q*_ is a diagonal matrix of *q*×*q* dimension. Residual deviations (***ε***) were assumed as ***ε*** ∼ *N*(***0***, ***I***_*n*_***σ***^***2***^), where *n* is the number of genotype-environment observations. Furthermore, the genotyping data were processed in two matrices of additive and dominance effects, modeled by:

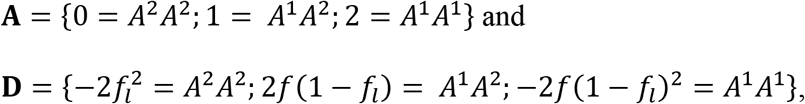

where *f*_*l*_ is the frequency of the favorable allele at locus *l*. Thus, the genomic-related kinships were estimated as follows:

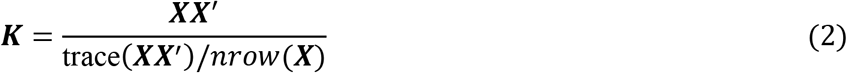

where ***K*** is a generic representation of the genomic kinship (***K***_A_, ***K***_D_), ***X*** is a generic representation of the molecular matrix (**A** or **D**), and *nrow*(***X***) denotes the number of rows in ***X*** matrix. Eq (2) was also used to shape the environmental relatedness kernels using T or W matrix. This linear kernel for **K**_*E*_ was described by Jarquín *et al*. (2014), which some other authors named it after “**Ω**”. Thus, here we only tested the difference between the enviromic source considered for building it and not the merit of the kernel method as was done in previous works (Costa-Neto *et al.,* 2021a).

#### 2.4.2 GBLUP with enviromic main effects from T matrix (E-GP)

From baseline equation (1), we include a main environmental relatedness effect carried out with the **T** matrix (***u***_E,T_), as follows (Costa-Neto *et al*., 2021a):

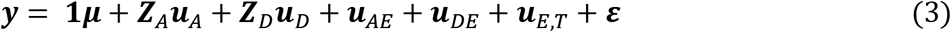

with 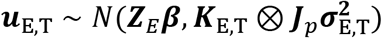, where ***J***_*q*_ is a *p*×*p* matrix of 1s, is ***K***_E,T_ the environmental relatedness created and variance component from the **T** matrix. If non-enviromic sources are considered, the expected value for environments is given by ***Z***_*E*_***β*** as the baseline model. In this model, the G×E effects are also modeled as the BD genomic matrix. Thus, we refer to this model as “E-GP (BD)”. The kernel of enviromic assembly (***K***_*E*,*T*_) was built using the panel of envirotype descriptors (**T**) in the same way as described in equation (2).

From model (3), we substitute the BD for a reaction-norm (RN, Jarquín et al., 2014) based on the Kronecker product between the enviromic and genomic kinships (Martini *et al*., 2020) for additive (***u***_*AE*,*T*_) and dominance effects (***u***_*DE*,*T*_):

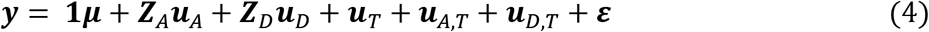

with 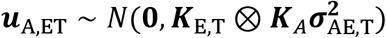 and 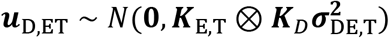 where 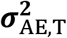 and 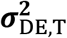 are the variance components for enviromic × additive and enviromic × dominance effects performed as reaction-norms (Costa-Neto *et al.,* 2021a; Rogers *et al*., 2021), respectively. For short, this model will be named “E-GP (RN)”.

#### 2.4.3 GBLUP with enviromic main effects from W matrix (W-GP)

Finally, in models (4) and (5), we substitute the enviromic assembly derived from **T** by the same kernel size derived from **W**, that is, an environmental relatedness with 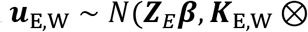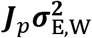), creating two more models:

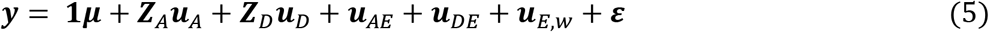

and

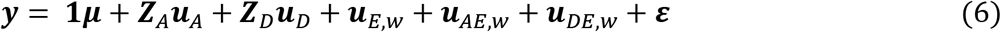

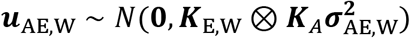 and 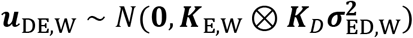, where 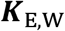 and 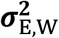 are the resulting kinship and the variance components estimated for enviromic assembly from the **W** matrix, respectively. Thus, for short, models (5) and (6) will be referred to as “W-GP (BD)” and “W-GP (RN)”, respectively.

### 2.5 Study cases for the E-GP platform

In this study, we conceived two cases to highlight the benefits of E-GP to boost efficiency in prediction-based platforms for hybrid development in maize breeding (Figure 2). The first case (*Case 1)* involves predicting the single-crosses from different theoretical existing experimental network setups, where we dissect the predictive ability over four prediction scenarios. In the second case (Case 2), we explore a theoretical conception of a super-optimized experimental network using the most representative combination of genotypes-environments selected using genomics, enviromic assembly, and genetic algorithms. Below we detail each case studied.

**Fig. 2.**
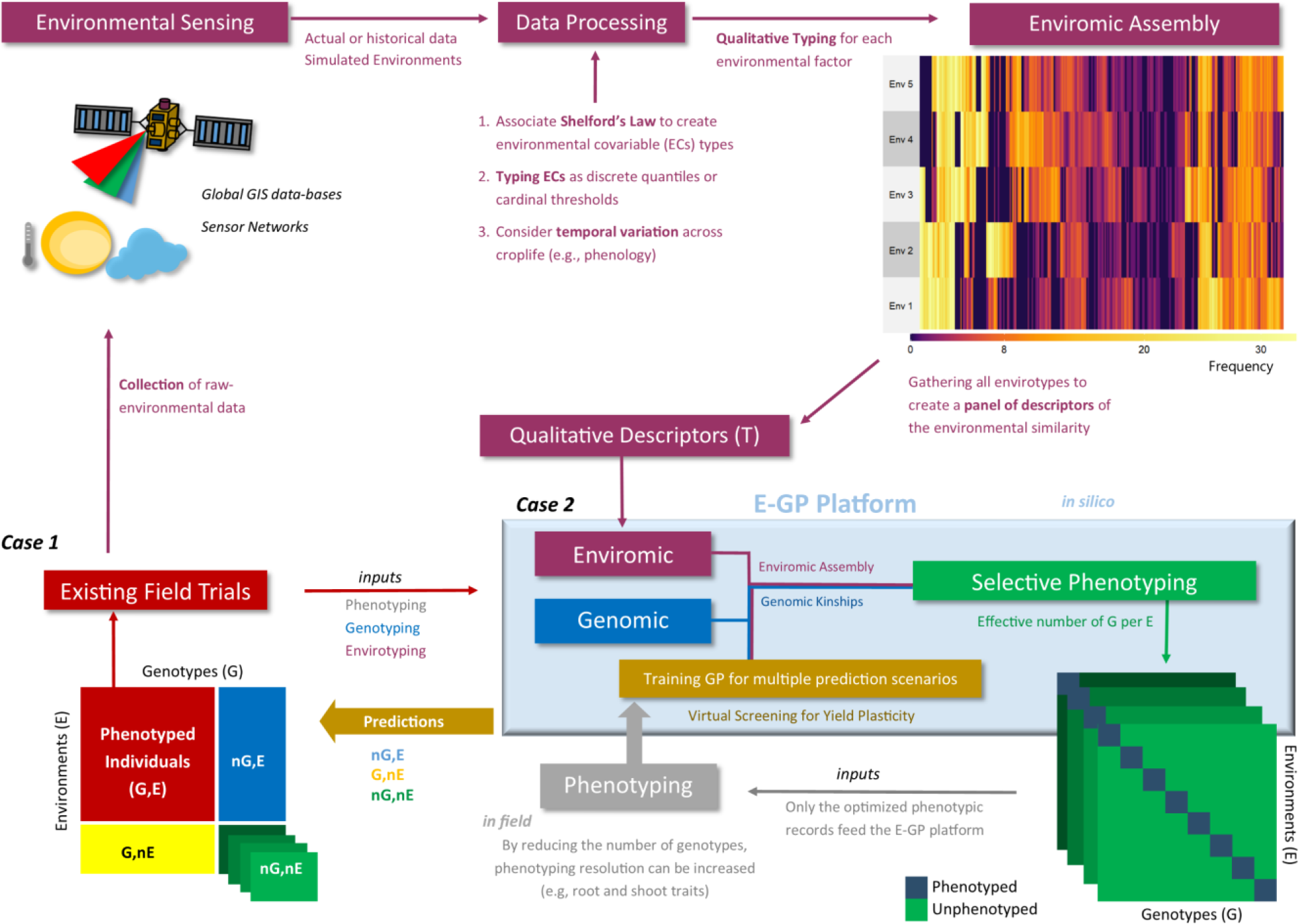
Workflow of the E-GP considering the two study cases (Case 1 and Case 2) of this study.

#### 2.5.1 Case 1: expanding the existing field trials

In the first case (*Case 1*), we design a novel cross-validation scheme to split the global available phenotypic information (*n*), from *p* genotypes and *q* environments, into different training setups. Consequently, four prediction scenarios were created based on the simultaneous sampling of the phenotypic information for *S* genotypes and *R* environments.

- G, E: refers to the predictions of the tested genotypes within the experimental network (known genotypes in known environmental conditions). The size of this set is 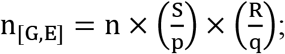
- *n*G,E: refers to predictions of untested (new) genotypes within the experimental network (known environmental conditions). The size of this set is 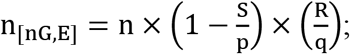
- G,*n*E: in this scenario, predictions are made under environmental conditions external to those found within the experimental network. However, there is phenotypic information available within the experimental network. The size of this set is 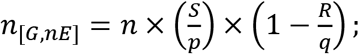
- *n*G,*n*E: refers to predicting untested (new) genotypes and untested (new) environmental conditions. This set’s size is 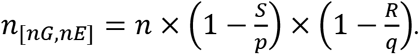

Theoretically, if *R*/*q* = 1, then *n*_[*G*,*nE*]_ = *n*_[*nG*,*nE*]_ = 0, equal to the commonly used CV1 scheme (prediction of novel genotypes in known environments). Different intensities of *R*/*q* can be sampled, which permits the testing of different sets of experimental networks. Here we simulated three different experimental network setups for each tropical maize data set. For the *N-level set*, we made 3/8, 5/8, and 7/8; for the *Multi-local set* 2/5, 3/5, and 4/5. We assumed the same level of genotype sampling as the training set for all experimental setups, equal to a fraction of 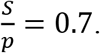 Each training setup was randomly sampled 50 times in order to compute the prediction quality statistics. For this purpose, two statistics were used to assess the statistical models’ performance over these training setups. We calculated Pearson’s moment correlation (*r*) between observed (*y*) and predicted (*y*^) values and used the average value for each model and training setup as a predictive ability statistic. To check the GP’s ability to replace field trials, we then computed the coincidence (CS, in %) between the field-based selection and the selection-based selection of the top 5% best-performing hybrids in each environment.

#### 2.5.2 Case 2: designing super-optimized field trials

The second case (*Case 2*) was performed on the optimized training set described below. The first step was to compute a full-entry G×E kernel, based on the Kronecker product (⊗) between the enviromic assembly-based relatedness kernel (***K***_*E*,*T*_, *q* × *q* environments) and genomic kinship (***K***_*G*_, *p* × *p* genotypes), thus***K***_*GE*,*T*_ = ***K***_*E*,*T*_ ⊗ ***K***_*G*_, with an *n* × *n* dimension, in which *n* = *pq*. Here we adopted the kernel made up for additive effects (***K***_*G*_ = ***K***_*A*_) as the genomic kinship, despite the benefits of dominance effects in modeling G×E. We chose to use only ***K***_*A*_ for simplicity and since additive effects seems to be a major genomic-related driver of G×E for grain yield in tropical maize (Dias *et al*., 2018; Alves *et al*., 2019; Costa-Neto *et al*., 2021a; Roger *et al*., 2021), a fact that was also observed for *Case 1* (see section 3.1). Later, we applied a single-value decomposition in ***K***_*GE*,*T*_, following ***K***_*GE*,*T*_ = ***UVU***^***I***^ where ***U*** is a total of eigenvalues and ***V*** the respective eigenvectors. The number of eigenvalues that explains 98% of the variance present in ***K***_*GE*,*T*_ indicate the number of effective SNPs by envirotype-marker interactions, which is also the minimum core of genotype-environment combinations (*N*_*GE*_). Thus, the reduced phenotypic information of some genotypes in some environments (*N*_*GE*_) was used to predict a virtual experimental network (*N*_*test*_), involving all remaining single-crosses in all available environments, thus given by *N*_*test*_ = *n* − *N*_*GE*_,

Following this step, a genetic algorithm scheme using the design criteria PEV_MEAN_ was used to identify the *N*_*GE*_ combinations of genotypes in environments within the ***K***_*GE*,*T*_ entries that must be phenotyped (Misztal, 2016). This optimization was implemented using the *SPTGA* R package (Akdemir and Isidro-Sánchez, 2019) using 100 iterations: five solutions selected as elite parents were used to generating the next set of solutions and mutations of 80% for each solution generated.

### 2.6 Virtual screening for yield plasticity

Finally, we tested each GP model’s potentials to predict the genotypes’ phenotypic plasticity and stability across environments using only the *N*_*GE*_ phenotypic information. First, the prediction ability was computed for genotypes by correlating the predicted and observed grain yield values across environments (Costa-Neto *et al*., 2021a). The second measure was based on the Finlay-Wilkinson adaptability model’s regression slope (Finlay and Wilkinson, 1963). The GP predicted values were regressed to the observed environmental deviations, as follows:

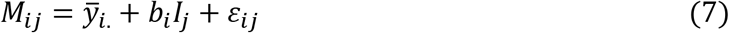

where *M*_*ij*_ is the expected GP-based mean value of grain yield for *i*^th^ genotype at *j*^th^ environment, *y̅*_*i*._ is the mean genotypic value for *i*^th^ genotype, *b*_*i*_ is the genotype plastic response across the mean-centered standardized environmental score (*I*_*j*_) and *ε*_*ij*_ is the variety of residual deviation sources not accounted in the model. After this step, the Pearson’s product-moment correlation between GP-based 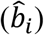 and phenotypic-enabled estimates were computed as an indicator of the ability to reproduce plastic responses *in silico* for the *p* genotypes. For this, the mean squared error is also calculated as:

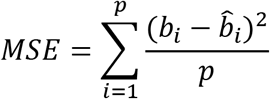

All statistics were computed using the entire data sets and only the top 5% of genotypes selected for each environment. The latter aimed to check the efficiency of the E-GP method to produce high-quality virtual screenings for plasticity.

### 2.7 Data and Code availability

All data sets and codes (in R) are freely available at https://github.com/gcostaneto/EGP.

## 3 RESULTS

### 3.1 Case 1: Accuracy in predicting diverse G×E scenarios

A cross-validation scheme was designed to assess the predictive ability of the enviromic-aided approaches in the face of traditional GBLUP. For that, sample genotypes (70%) and environments were used to compose a drastically sparse training set for MET (training environments/total of environments). This helped assess the efficiency of E-GP for *Case 1*, in which we were able to dissect the predictive ability (section 3.2.3) in different scenarios of a scarcity of phenotypic records: novel genotypes in tested environments (*n*G,E); tested genotypes in untested environments (G,*n*E), and novel genotype and environment conditions (*n*G, *n*E). Tables 1 and 2 present the N-level and Multi-Regional sets results, respectively. Then, these results were gathered for both data sets and four prediction scenarios in order to check for the joint predictive ability analysis (Figure 3).

**Fig. 3.**
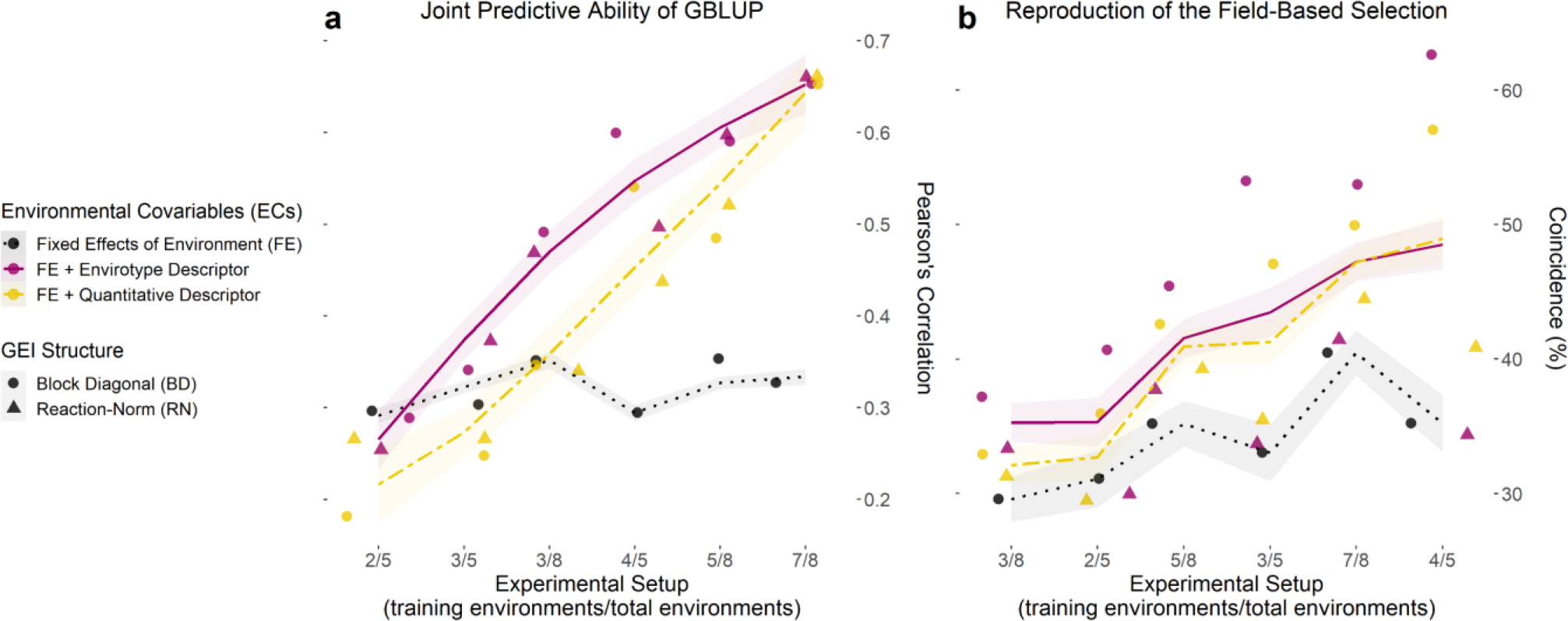
Joint accuracy trends of GP models for each training setup of existing experimental networks. **a**. Predictive ability computed with the correlation (*r*) between observed (*y*) and predicted (*ŷ*) values for the grain yield of each genotype in each environment, over three experimental setups (number of environments used/total of environments) for both maize sets (N-level and Multi-local), using 70% of the genotypes as a training set and the remaining 30% as a testing set. **b**. Coincidence index (CS) between the field-based and prediction-based selection of the best 5% genotypes in each environment for the same experimental setups and data sets. Dots and triangles represent the point estimates of predictive ability and CS for models involving a block diagonal genomic matrix for G×E effects (dotted) and an enviromic × genomic reaction-norm G×E effect (triangle). Trend lines were plotted from the partial values of each sample (from 1 to 50) and three prediction scenarios (*n*G, E; G, *n*E and *n*G, *n*E) by using the gam() integrated with smoothness estimation in R. Black dotted lines represent the benchmark GBLUP method, considering the effect of the environment as a fixed intercept. Yellow two-dash lines represent the GBLUP involving the main effect from quantitative descriptors (**W** matrix). Finally, solid dark pink lines represent the GBLUP involving the main effect of envirotype descriptors (**T** matrix). Thus, the latter represents the E-GP based approach for *Case 1* (predictions under existing experimental networks).

**Table 1.** Predictive ability (± standard error) of the genome-based prediction models (GP) for the N-level set of tropical maize hybrids (570 hybrids × 2 locations × two years × two nitrogen managements). Bold values denote higher predictive ability values for each scenario: G,E (known genotypes at known growing conditions), G,*n*E (known genotypes at new growing conditions), *n*G, E (new genotypes at known growing conditions), and *n*G, *n*E (new genotypes at new growing conditions).

| Training Setup | Model | Prediction Scenario |  |  |  |
| --- | --- | --- | --- | --- | --- |
|  |  | G, E | G, nE | nG, E | nG, nE |
| 7/8<br>Environments | GBLUP | 0.771± 0.064 | 0.397±0.046 | 0.310±0.054 | 0.297±0.029 |
| | E-GP (BD) | **0.903**±0.115 | **0.493**±0.169 | **0.615**±0.022 | **0.416**±0.153 |
| | E-GP (RN) | 0.833±0.118 | **0.477**±0.199 | **0.613**±0.040 | 0.394±0.193 |
| | W-GP (BD) | **0.915**±0.115 | 0.333±0.208 | **0.614**±0.025 | 0.242±0.189 |
| | W-GP (RN) | 0.885±0.117 | 0.327±0.210 | 0.613±0.031 | 0.23±0.196 |
| 5/8<br>Environments | GBLUP | 0.747±0.049 | 0.432±0.046 | 0.294±0.026 | 0.323±0.04 |
| | E-GP (BD) | 0.905±0.056 | **0.554**±0.144 | **0.659**±0.015 | **0.464**±0.113 |
| | E-GP (RN) | 0.833±0.056 | **0.570**±0.132 | **0.660**±0.025 | **0.475**±0.104 |
| | W-GP (BD) | **0.931**±0.057 | 0.449±0.286 | 0.659±0.019 | 0.347±0.253 |
| | W-GP (RN) | 0.897±0.056 | 0.501±0.229 | **0.660**±0.026 | 0.395±0.198 |
| 3/8<br>Environments | GBLUP | 0.739±0.040 | 0.527±0.080 | 0.295±0.015 | 0.394±0.044 |
| | E-GP (BD) | 0.899±0.026 | 0.534±0.081 | 0.660±0.012 | 0.388±0.038 |
| | E-GP (RN) | 0.823±0.026 | **0.566**±0.086 | **0.663**±0.015 | **0.420**±0.041 |
| | W-GP (BD) | **0.924**±0.026 | 0.532±0.08 | 0.660±0.015 | 0.384±0.038 |
| | W-GP (RN) | 0.886±0.025 | **0.579**±0.088 | **0.663**±0.020 | **0.424**±0.041 |

**Table 2.**
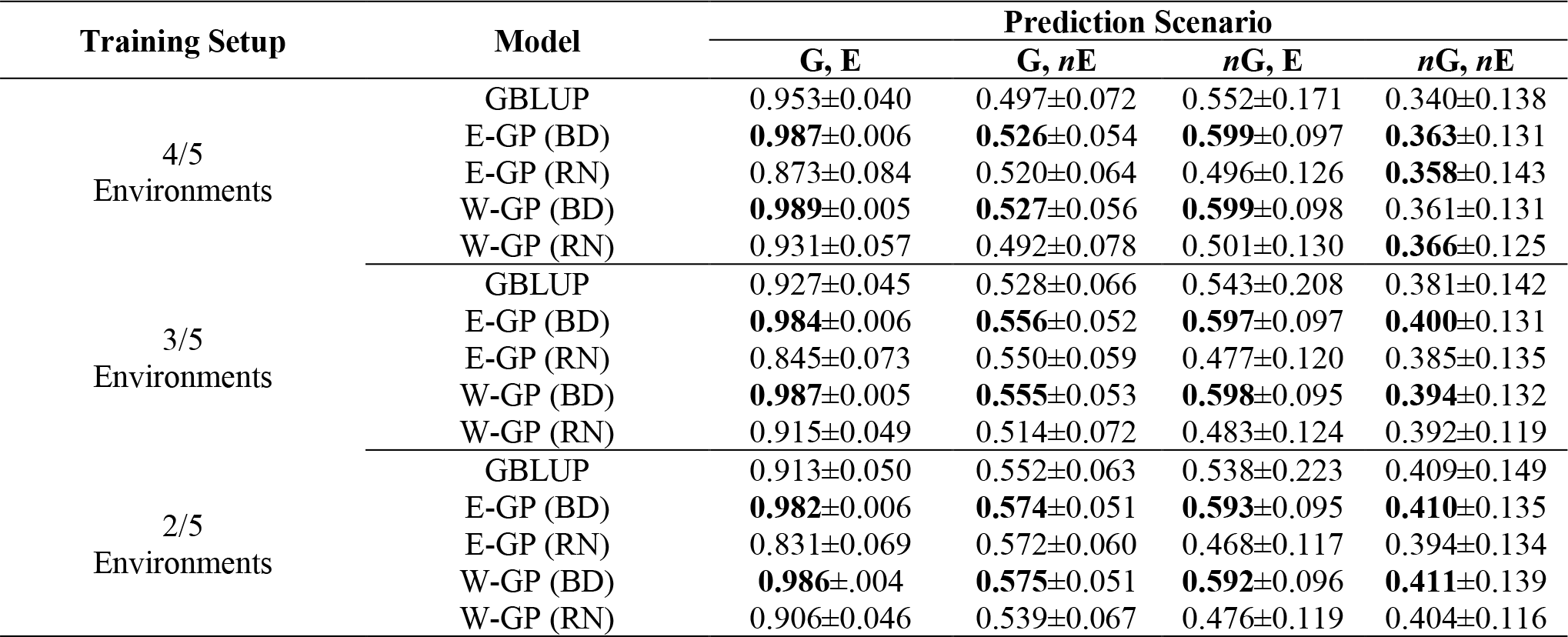
Predictive ability (± standard error) of the genome-based prediction models (GP) for the Multi-Local set of tropical maize hybrids (247 hybrids × 5 locations in different regions of Brazil). Bold values denote the higher predictive ability values for each scenario: G,E (known genotypes at known growing conditions), G,*n*E (known genotypes at new growing conditions), *n*G, E (new genotypes at known growing conditions), and *n*G, *n*E (new genotypes at new growing conditions).

| Training Setup | Model | Prediction Scenario |  |  |  |
| --- | --- | --- | --- | --- | --- |
|  |  | G, E | G, nE | nG, E | nG, nE |
| 4/5<br>Environments | GBLUP | 0.953±0.040 | 0.497±0.072 | 0.552±0.171 | 0.340±0.138 |
| | E-GP (BD) | **0.987**±0.006 | **0.526**±0.054 | **0.599**±0.097 | **0.363**±0.131 |
| | E-GP (RN) | 0.873±0.084 | 0.520±0.064 | 0.496±0.126 | **0.358**±0.143 |
| | W-GP (BD) | **0.989**±0.005 | **0.527**±0.056 | **0.599**±0.098 | 0.361±0.131 |
| | W-GP (RN) | 0.931±0.057 | 0.492±0.078 | 0.501±0.130 | **0.366**±0.125 |
| 3/5<br>Environments | GBLUP | 0.927±0.045 | 0.528±0.066 | 0.543±0.208 | 0.381±0.142 |
| | E-GP (BD) | **0.984**±0.006 | **0.556**±0.052 | **0.597**±0.097 | **0.400**±0.131 |
| | E-GP (RN) | 0.845±0.073 | 0.550±0.059 | 0.477±0.120 | 0.385±0.135 |
| | W-GP (BD) | **0.987**±0.005 | **0.555**±0.053 | **0.598**±0.095 | **0.394**±0.132 |
| | W-GP (RN) | 0.915±0.049 | 0.514±0.072 | 0.483±0.124 | 0.392±0.119 |
| 2/5<br>Environments | GBLUP | 0.913±0.050 | 0.552±0.063 | 0.538±0.223 | 0.409±0.149 |
| | E-GP (BD) | **0.982**±0.006 | **0.574**±0.051 | **0.593**±0.095 | **0.410**±0.135 |
| | E-GP (RN) | 0.831±0.069 | 0.572±0.060 | 0.468±0.117 | 0.394±0.134 |
| | W-GP (BD) | **0.986**±0.004 | **0.575**±0.051 | **0.592**±0.096 | **0.411**±0.139 |
| | W-GP (RN) | 0.906±0.046 | 0.539±0.067 | 0.476±0.119 | 0.404±0.116 |

#### 3.1.1 Within experimental network (know growing conditions)

The predictions within known environmental conditions of a certain experimental network involve scenarios *G,E* and *nG,E*. For the *G,E* scenario (classical ‘training set’), all models outperformed the GBLUP in any setups N-level set, and most of the setups of Multi-Regional set. The highest values of predictive ability were observed for enviromic-aided GP models using the block-diagonal matrix for G×E effects (BD), that is, the E-GP (BD) and W-GP (BD), respectively. Two general trends were observed: the size of the experimental setup has a small effect on GP models’ accuracy. Secondly, higher accuracy gains were observed for the N-level set (Table 1), with a higher number of entries (more genotypes and more environments). The accuracy gains in this N level set ranged from +8% (*r* = 0.83 for E-GP RN at 7/8 experimental setup), in relation to *r* = 0.77 (GBLUP), to +24% (*r* = 0.92 for W-GP RN at 3/8 experimental setup), in relation to *r* = 0.74 (GBLUP). In contrast, for the Multi-Regional set (Table 2), both RN-G×E models reduced the accuracy (on average, -3%). For the BD-G×E models, small gains in accuracy (from +4% to +8%) were observed.

That is also a trend for the second prediction scenario (*n*G,E), in which the Multi-Regional set presented an average gain of 10% for all enviromic-aided GP models with BD-G×E, and a reduction of 10% for all RN-G×E models. Conversely to the previous scenario (G,E, within the experimental network, using known genotypes), the *n*G,E is one of the most important plant breeding scenarios. It represents the ability of predict new single-crosses, within the know environmental gradient, borrowing genomic and enviromic information from the phenotypes of the relatives. Thus, expand the spectrum of possible genotypes using know growing conditions from the past. For the N-level set, gains up to 100% were observed for all enviromic-aided models using any G×E structure. No differences were observed between enviromic-aided models and experimental setups. On average, all enviromic-aided models achieved a predictive ability of approximately *r* = 0.66 across all experimental setups (3/8, 5/8, and 7/8, Table 1). In contrast, the GBLUP model has been impacted with reduced accuracy and a lack of phenotypic records. The highest gains in predictive ability were observed for scenario 3/8, average +118% for BD-G×E models, and +119% for RN-G×E models.

#### 3.1.2 Across experimental network (new growing conditions)

The predictions within new environmental conditions across the experimental network involve *G,nE* and *nG,nE*. Both scenarios represent the ability of using the available phenotype information collected from experimental network in order to predict novel growing conditions using genomic or genomic+enviromic data sources. For the *G,nE*, the E-GP models outperformed W-GP and GBLUP across most experimental setups, despite small differences between the enviromic-aided approaches. For the E-GP BD at the N-level set (Table 1), the gains in predictive ability ranged from +24% (*r* = 0.49 at 7/8 setup, Table 1), in relation to *r* = 0.40 (GBLUP), to +35% (*r* = 0.57 at 5/8 setup), in relation to *r* = 0.43 (GBLUP). However, for scenario 3/8, these gains were equal to +10% (*r* = 0.57) in relation to the +13% archived by the benchmark W-GP RN (*r* = 0.58), both over the *r* = 0.53 from GBLUP. In scenario 7/8, W-GP was outperformed by GBLUP, with a reduction in accuracy between -18% and -16%, where the E-GP made better use of the large phenotypic information available for training GP models (gains from +20% to +24% over GBLUP). A similar pattern was observed for the Multi-Regional set (Table 2), in which the gains of E-GP ranged from +4% to +6% across all setups, and W-GP ranged from -3% to +6% under the same conditions.

The second scenario involving novel growing conditions also predicts novel genotypes (*nG,nE*) into account. Thus, all predictions were based on the phenotypic records from reassembled genotypes and considering the environmental similarity conceived from enviromics. With a large experimental network and genomics, the E-GP models outperformed W-GP and GBLUP when predicting new G×E. Observed accuracy gains ranged from 33% (*r* = 0.39 for E-GP RN) to 40% (*r* = 0.42 for E-GP BD), in experimental setup 7/8 (Table 1), where GBLUP achieved *r* = 0.30, and from 47% (*r* = 0.46 for E-GP BD) to 51% (*r* = 0.48 for E-GP BD), at the experimental setup 5/8, where GBLUP achieved *r* = 0.32. Unlike observations in the other prediction scenarios, the models RN-G×E outperformed BD-G×E in experimental setups 3/8 (N-Level set) and 2/5 (Multi-Regional set).

### 3.2 Accuracy trends across diverse experimental setups

This section highlights the main target of our *Case 1* study, in which the predictive ability was achieved using the merged information of scarce genotypes at some environments. Joint accuracy trends showed that E-GP was useful at increasing GP accuracy (Fig. 3a) and explaining the phenotypic variation sources in both maize data sets (Supplementary Table 3-4). For scenarios with reduced phenotypic information (e.g., 3/5, 3/8, and 4/8), any model with some degree of environmental information outperformed the GBLUP for all scenarios. The E-GP approach (purple colors in Figure 3a) better captured envirotype-phenotype relations and converted them into accuracy gains among these models. This is also reflected in the E-GP efficiency as a predictive breeding tool capable of reproducing field-based trials (Fig. 3b).

Regarding the G×E structures, the contribution of RN-G×E is significant only for drastically lacking phenotypic records (training setup 3/8), leading to the conclusion that the use of a main-effect is substantial for most cases E-GP is enough to increase accuracy in GBLUP. For setup 2/5 (Multi-Regional Set), no differences were observed between all the GP models.

The coincidence between the GP-based selection and the in-field selection (CS, %) ranged from ∼35% to ∼50%, in models with some environmental information, while it ranged between 30% and 40% for GBLUP (without environmental information). For the E-GP approach accounting for a wide number of phenotypic records in the training set (7/8, 3/5, and 4/5), values of CS up to 55% were found. Among these models, it seems that the RN-G×E reduces the CS estimates concerning the BD-G×E based models. Considering both figures 3a and 3b, it is possible to suggest that predictive ability does not imply an increase of CS, that is, in the power of selecting the best performing genotypes in certain environments. However, the drastic increase in the E-GP accuracy in relation to the other models leads us to infer that despite the lower rise in CS, the E-GP models are useful when predicting GE for a vast number of single-crosses.

### 3.3 Case 2: enviromic assembly with optimized training sets for genomic prediction

Those results lead us to investigate *Case 2* (Fig. 2), where we checked the possibility of training efficient and biologically accurate GP scenarios from super-optimized training sets. Then, we checked the potential of using these optimized field trials for predicting novel G×E under the so-called “virtual experimental networks”. This approach were implemented by combining two selective phenotyping approaches (Misztal, 2016; Akdemir and Isidro-Sánchez, 2019), aiming to identify combinations of genotypes and environments using in-silico representations of the enviromic assembly × genomic kinships.

#### 3.3.1 Predicting G×E at virtual experimental networks

The process of designing virtual networks in maize hybrid breeding involved two steps (Supplementary Fig 1). First, we used a single-value decomposition (SVD)-based algorithm to select the effective number of individuals (*N*_GE_) (Misztal, 2016) representing at least 98% of the variation of ***K***_*G*,*ET*_. It was done in ***K***_*G*,*ET*_ because this kernel represents an in-silico representation of envirotypes and genotypes (Akdemir and Isidro-Sánchez, 2019). Under sparse MET conditions, it led to a training size equal to *N*_GE_ = 67 and *N*_GE_ = 49 for the N-level set (*n* = 4,560) and Multi-Regional set (*n* = 1,235), respectively. It represents only 1.5% and 4% of the whole experimental network; Supplementary Fig. 2-3. For didactic purposes, from here onwards, we will represent the values of *N*_GE_ as the training set size/number of genotypes.

We also checked the use of all environments, although the accuracy differences were tiny in relation to this sparse MET scenario (Table 3). Furthermore, small differences were achieved by E-GP and W-GP models with BD-G×E, but both higher than RN-G×E and GBLUP (Fig 4). Major differences were highlighted as follows:

- For within-field trials, predictive ability ranged from *r* = 0.76 (W-GP) to *r* = 0.87 (E-GP);
- For virtual-networks, it ranged from *r* = 0.14±0.11 (GBLUP) to *r* = 0.60±0.06 (E-GP);
- In virtual-networks, the predictive ability of models trained with drastically reduced phenotypic records ranged from *r* = 0.10 (GBLUP, N_GE_ = 67/4560) to *r* = 0.58 (E-GP, N_GE_ = 67/4560) and *r*=0.18 (GBLUP, N_GE_ = 49/1235) to *r*=0.81 (E-GP, N_GE_ = 49/1235).

**Fig. 4.**
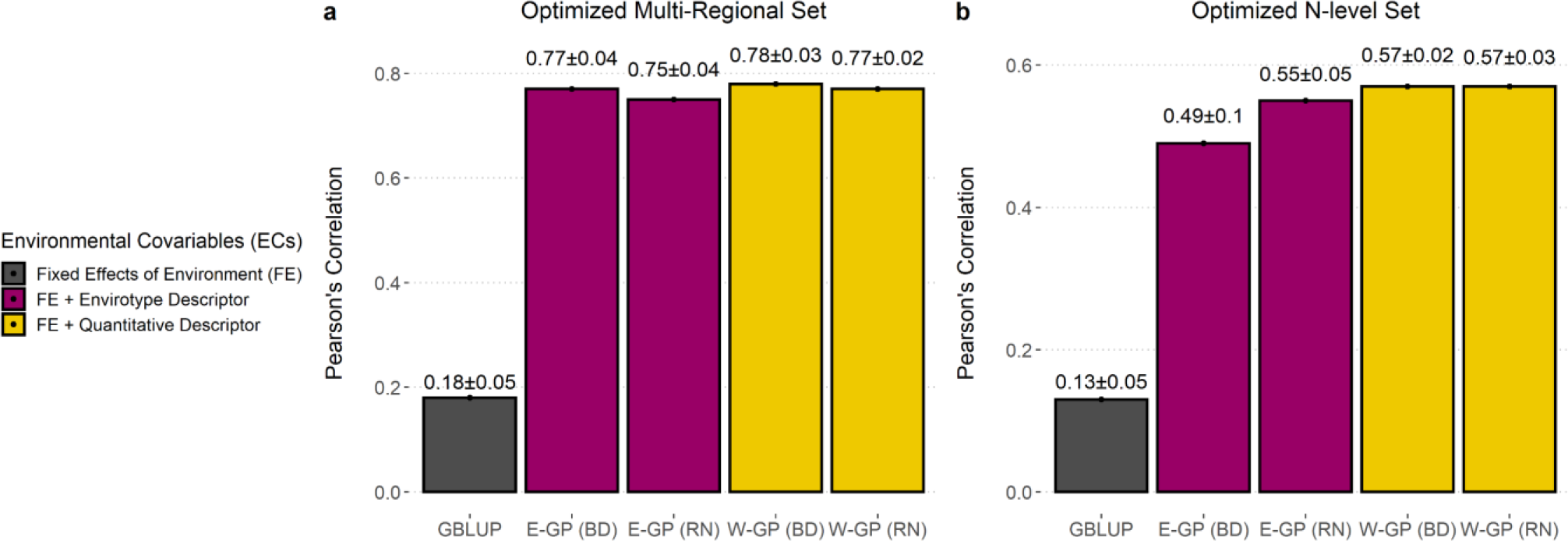
Accuracy of GP models trained with super-optimized experimental networks. Predictive ability (*r*) plus standard deviation measured by the correlation between observed and predicted values for each model in the optimized Multi-Regional Set (**a**); and for the N level Set (**b**). Barplots were colored according to the type of environmental covariable (ECs) used: none (black), envirotype descriptor (**T** matrix, wine), and quantitative descriptor (**W** matrix, yellow).

**Table 3.**
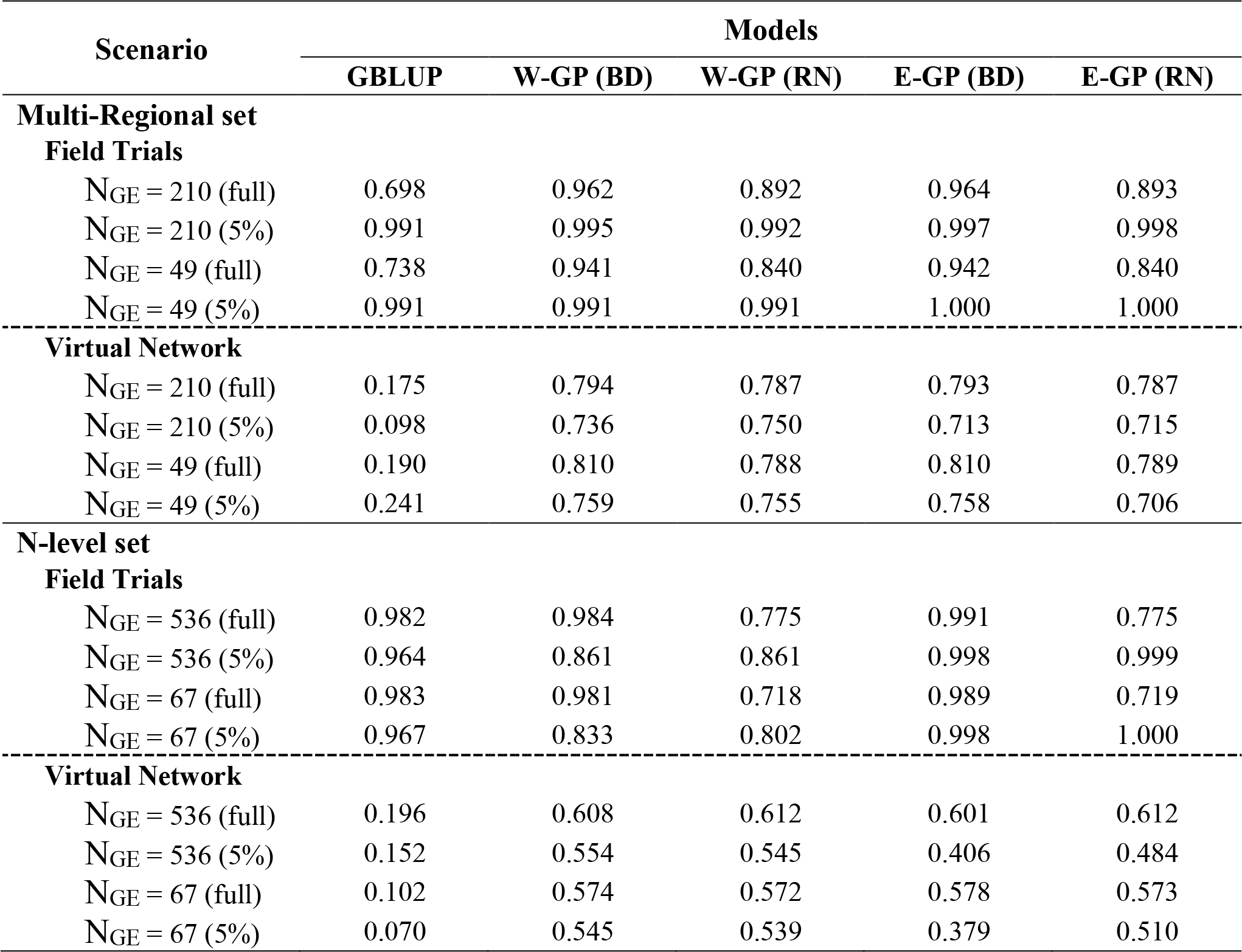
Predictive ability of the genomic prediction models (GP) for two tropical maize data sets (Multi-Regional and N-level) produced using the effective number of phenotypic records (N_GE_, genotypes-environments observations) and for the scenarios Field Trials (predicting N_GE_) and Virtual Network (predicting *n* – N_GE_, where n is the number of genotypes by environments available in the full data set). The reference “full” and “5%” in parentheses represents the predictive ability produced with all genotypes and using only the top 5%, respectively

The predictive ability was computed considering only the top 5% of genotypes in each environment and data set. The objective was to verify if the GP approaches could adequately predict the performance of the best-evaluated genotypes in the field. For the Multi-Regional set, the predictive ability ranged from *r* = 0.098 (GBLUP, N_GE_ = 210/1235) to *r* = 0.579 (W-GP BD, N_GE_ = 49/1235) and *r* = 0.578 (E-GP BD, N_GE_ = 49/1235); For the N-level set, W-GP outperformed E-GP, leading to *r* = 0.554 (W-GP BD, N_GE_ = 536/4560) in front of *r* = 0.554 (E-GP RN, N_GE_ = 67/4560) but with less phenotyping data. In contrast, the best E-GP model at the higher number of genotypes and environments evaluated in the field *r* = 0.484 (E-GP RN, N_GE_ = 536/4560) were outperformed by the same model, yet with less phenotyping data *r* = 0.554 (E-GP RN, N_GE_ = 67/4560). For GBLUP, the effective size of the training set was important, ranging in predictive ability from *r* = 0.070 (N_GE_ = 67/4560) to *r* = 0.152 (N_GE_ = 536/4560). The result of both sets suggests that when using enviromic-aided approaches, the use of fewer amounts of, but more representative, phenotyping information is better than more amounts of, yet less representative, phenotyping data.

Figure 4 was created using the average values of Table 3. This figure shows that the optimization was more effective for growing conditions contrasting across macro-regions (Fig. 4a) than for experimental networks involving fewer locations (Fig. 4b). Notably, it is possible to drastically reduce field costs for experimental networks conducted across diverse locations. However, for screening management conditions, greater precautions must be considered with the use of E-GP.

#### 3.3.2 Predicting genotype-specific plasticity and environmental quality

In this step, we checked these models’ ability to produce virtual screenings for yield plasticity (Fig.5). We used the Finlay-Wilkinson method (FW, Eq. 7) over the predicted GY means of each genotype *i* in environment *j* (*M*_*ij*_). Hence, we compared the ability of E-GP in the prediction of: (1) individual genotypic responses across environments, (2) the gradient of environmental quality (ℎ_*j*_), and (3) the plasticity coefficient (*b*_1_) of the FW model describing the rate of responsiveness to *h*. The results in Fig 5 involves a joint analysis of both data sets.

**Fig. 5.**
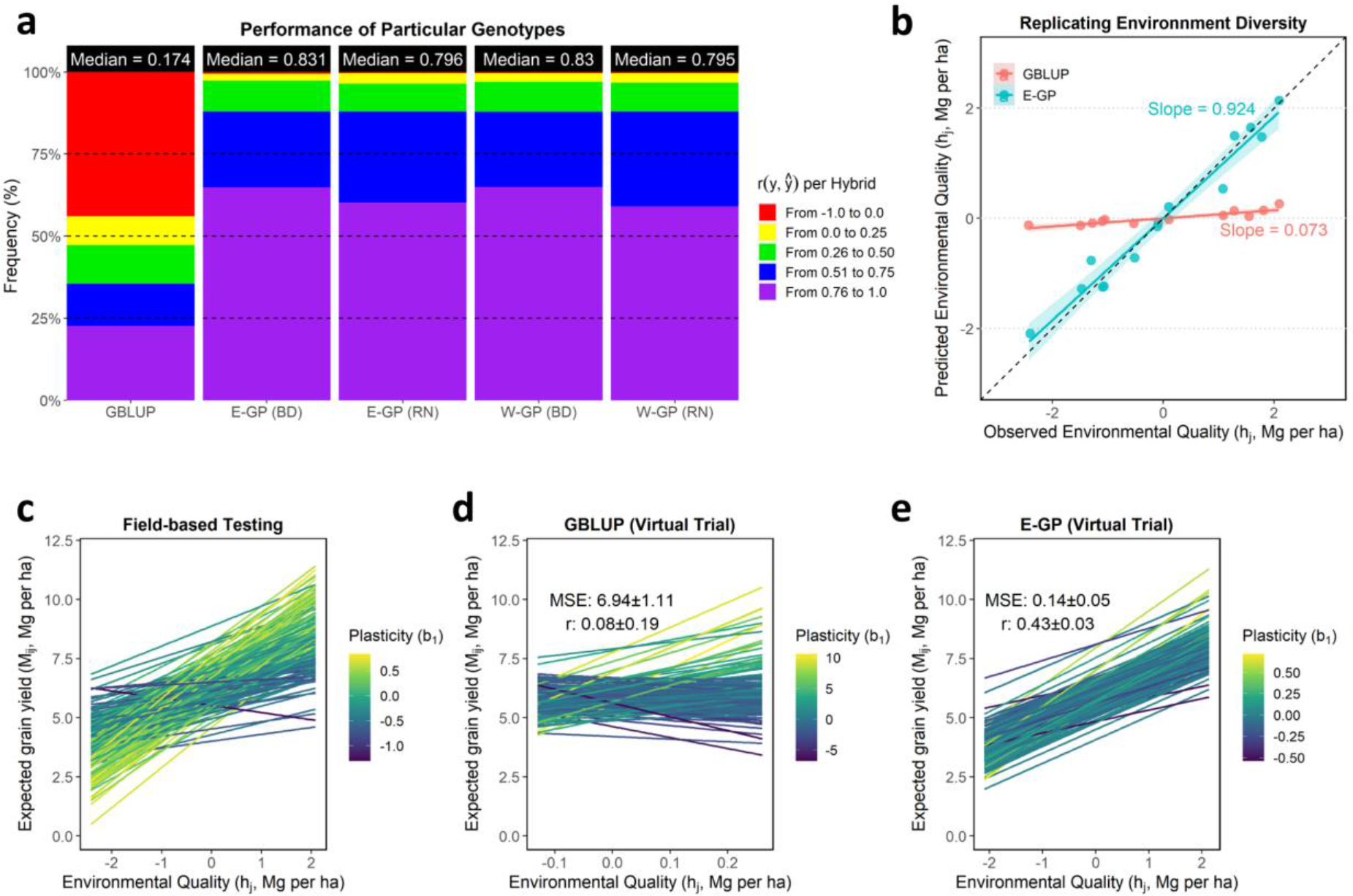
Accuracy of GP models in reproducing the genotype-specific plasticity. **a**. The panel of predictive ability (*r*) explaining the plasticity of genotypes across environments. This statistic was estimated for each individual (hybrid) by correlating observed and predicted values across environments. Individuals with values below 0 were considered unpredictable and marked in red. **b**. ability of the prediction-based tools to reproduce an existing experimental network’s environmental quality (*h_j_*). In the X-axis, we find the *h_j_* computed using the phenotypic records of a current experimental network. In the Y-axis, the *h*_j_ values are presented considering a virtual experimental network built up using GBLUP and E-GP (with BD) predictions. **c**-**e**. Yield plasticity panels denoting each genotype’s G×E effects across the *h_j_* values for observed field-testing screening (**c**) concerning prediction-based (**d**-**e**). Only the 5% best genotypes in each environment were used to create this plot. Each line was colored with the genotype-specific plasticity coefficient (*b*_1_). For the N-level set, the full-optimized set (536 hybrids over eight environments) was used.

All models that included some degree of enviromic assembly outperformed the GBLUP-based approach when predicting individual genotype responses across the MET (Fig 5a). The median values of *r* ranged from *r* =0.17 (GBLUP), in which 45% of the genotypes were not well predicted (red colors), to *r* = 0.83 (E-GP), in which up to 60% of the genotypes were very well predicted (purple colors). The inclusion of any enviromic assembly and G×E structure led to drastic gains in accuracy for a particular genotype response across contrasting (and unknown) G×E conditions (gains up to ∼378%). However, the BD structure outperformed RN in terms of resolution (many purple colors in Fig 5a). A major part of the accurately predicted performance of genotypes across environments ranged from *r* = 0.75 to *r* =1.0. Due to this, for the next figures, we plotted only the E-GP considering the BD-G×E structure.

GBLUP was unable to correctly reproduce ℎ_*j*_ for an in-silico study using the FW model (Fig. 5b). We observe that E-GP better describes the ℎ_*j*_ gradient (mean-centered average values of GY for each environment), with *r* near to 1 (correlation between observed and predicted environmental quality) also suggesting a low bias (*slope* = 0.924 between observed and predicted values). Consequently, this was reflected in the quality of yield plasticity predictions (Fig. 5c-e), as yield plasticity was represented as linear responsiveness over the environmental variation. The graphical representation of genotype-specific linear reaction norms dictated by the linear regression slope (*b*_1_) was likely more similar to E-GP than GBLUP about those observed in field-based testing (Fig 5b). The accuracy for *b*_1_ ranged from *r* = 0.08 (GBLUP) to *r* = 0.43 (E-GP), an increase of 437%.

## 4 DISCUSSION

Large-scale envirotyping, or simply enviromics, is an emerging field of data science in agricultural research and modern breeding program routines. We demostrated that enviromics is the science capable of bringing together environment information and quantitative genomics into an ecophysiology-smart manner. In this study, we presented the first report on (1) the use of Shelford’s Law to guide the assembly of the enviromics for predictive breeding purposes over experimental networks; (2) the integration of enviromic assembly-based kernels with genomic kinship into optimization algorithms capable of designing selective phenotyping strategies and (3) a break of the paradigm relying on the fact that phenotyping a higher number of genotypes at higher number of environments do not always contribute to increasing the accuracy of GP for contrasting G×E scenarios, but there are pieces of evidences suggesting that enviromics increases accuracy under sparse multi-environment networks; (4) report that the process of deriving markers of environmental relatedness, here named ‘enviromic assembly’, is crucial for the implementation of low-cost GP platforms over multi-environmental conditions.

In this study, we also envisage that the process of enviromic assembly is supported by a strong theoretical background in ecophysiology, illustrating the potential uses of environmental information to increase the accuracy of predictive breeding for yield and plasticity. Our results indicate that the E-GP platform (Figure 2) can fit two types of prediction scenarios in plant breeding programs: (1) better use of the available phenotypic records to train more accurate GP models capable of aiding the selection of genotypes across multi-environmental conditions and (2) a method that reduces costs for field-based testing and enables an early screening for yield plasticity under crossover G×E conditions. Furthermore, we show that any model with some degree of enviromic assembly (by typology or quantitative descriptors) is always better to reproduce the genotypes’ environmental quality of field trials and phenotypic plasticity.

Below we discuss the aspects that support the use of E-GP for multi-environment predictions, involving the importance of breaking the paradigm that states that enviromics are not necessary to predict G×E accurately. We then discuss how the genomic and enviromic sources are linked in the phenotypic records collected from the fields and how this type of knowledge can improve the quality of the prediction-based pipelines for crop improvement. Finally, we envisage possible environmental-assembly applications supporting other predictive breeding fields, such as optimizing crop modeling calibration and how it can couple a novel level of climate-smart solutions for crop improvement as anticipating the plasticity of a large number of genotypes using reduced phenotypic data.

### 4.1 Why are enviromics important for multi-environment genomic prediction?

Genomic prediction (GP) platforms were first designed to model the *genotype-to-phenotype* relations under single environment conditions, e.g., in a breeding program nursery (Lorenzana and Bernardo, 2009; Windhausen *et al*., 2012; Zhao *et al*., 2012; Zhang *et al*., 2015). Under these conditions, the micro-environmental variations within breeding trials (e.g., spatial gradients in soil properties) are minimized in the phenotypic correction step by separating useful genetic patterns and experimental noises (non-genetic patterns). However, those phenotypic records carry the indissoluble effects of macro-environmental fluctuations of certain weather and soil factors that occurred during crop growth and development (Li *et al*., 2018; Vidotti *et al*., 2019; Millet *et al*., 2019; Guo *et al*., 2020; Jarquín *et al*., 2020). That seems to be of no concern when predicting novel genotypes under these same growth conditions (the CV1 scheme for single-environment models) yet becomes noise for multi-environment prediction scenarios. It is a consequence of the macro-environment fluctuations in the lifetime of the crops (Allard and Bradshaw, 1964; Bradshaw, 1965; Arnold *et al*., 2019), responsible for modulating the rate of gene expression (e.g., Jończyk *et al*., 2017; Liu *et al*., 2020) and fine-tuning epigenetic variations and related to transcriptional responses (e.g., Vendramin *et al*., 2020; Cimen *et al*., 2021).

For each unit that we call “environment” (field trial at the specific year, location, planting date, and crop management), there are various environmental factors such as water availability, canopy temperature, global solar radiation, and nutrient content in the soil. The expression of some genotype in some phenotype is then limited by the certain key environmental factors, acting in different levels of crop development as preconized by School of de Wit’ since 1965 (see Bouman *et al*., 1996). However, we revisited the Shelford’s theory, which suggests that a population’s fitness is given by the amount and distribution of resources available for its establishment and adaptation (Shelford, 1931). Thus, we reinterpret this concept by assuming that the relation between input availability (deficit, optimum amount, or excess) across different crop development stages drives the amount of the genetic potential expressed in phenotypes produced by the same genotype for a given environment. Therefore, it provides the foundations to elaborate the argument that there is also an indissoluble *envirotype-phenotype covariance* in the phenotypic records that is interpreted as a G×E interaction for each environment. Because of that, we envisage that any environmental relatedness kernel must account for it in any way.

The pioneer approaches to measuring crop adaptability use the average value of a given trait in a given environment as an environmental quality index (e.g., Finlay and Wilkinson, 1963). However, the problem with this approach is that it explains the quality of the environment realized by the genotypes evaluated in it, making it inefficient to explain the drivers of environmental quality and incapable of predicting untested growing conditions, as observed in our results for *Case 2* using GBLUP without enviromic data. In addition, our results for *Case 1* highlight that it is a limit in accuracy for traditional GBLUP across MET, in which the accuracy remains almost the same, regardless of the number of phenotypic records available.

A second intrinsic covariance can interpret this last result within the phenotypic records, which is the *genotype-envirotype covariance*. By adapting the Quantitative Genetics theory to the terminology used here, we can infer that each genotype reacts differently to each envirotype, resulting in a given phenotype. This phenotype is then used to provide small crop phenology differences (genetically determined window sizes for each development stage). Recent but pioneer works have been carried out to understand the genetic and environmental determinants of flowering time in sorghum (Li *et al*., 2018) and rice (Guo *et al*., 2020). That can be indirectly interpreted as cardinal differential thresholds for temperature response. Jarquín *et al* (2020) proved that it is possible to increase the ability of GP in predictive novel G×E by coupling information of day-length in the benchmark GP models. For all these examples reported above, we can infer that, when trying to predict a novel genotype, by borrowing genotypic information from the relatives at different environments, it is impossible to reproduce the genotype-envirotype covariance without adding any enviromic information into the model.

The presence of both *genotype-envirotype* and *envirotype-phenotype* covariances might explain the gains in the predictive ability due to the use of multi-environment GP models in contrast to single-environment GP models (Bandeira e Souza *et al*., 2017; de Oliveira *et al*., 2020) and why deep learning approaches have successfully captured intrinsic G×E patterns and translated them into gains in accuracy (Montesinos-López *et al*., 2018; Crossa *et al*., 2019; Cuevas *et al*., 2019). Conversely, this also might explain the need to incorporate secondary sources of information in the prediction of grain yields across multiple environments (Westhues *et al.,* 2017; Ly *et al.,* 2018; Millet *et al.,* 2019; Costa-Neto *et al*., 2021a; 2021b; Jarquín *et al*., 2020), as well as the possible limitations of CGM approaches contrasting scenarios differing from those targeted near-iso conditions of CGM calibration (e.g., Cooper *et al*., 2016; Messina *et al*., 2018). Thus, an alternative can be supervised approaches to describe the environmental relatedness, such as in this paper, and perhaps unsupervised algorithms capable of taking advantage of the covariances related to the genotype-phenotype, genotype-envirotype, and envirotype-phenotype dynamics.

### 4.2 Sometimes main-effect enviromics is better than reaction-norm models

Our results from *Case 1* show that the inclusion of enviromic sources (for main-effects or explicitly incorporated in the RN-G×E structure) led to a better description of the envirotype-phenotype covariances, which was reflected in accuracy gains. At our data and Bayesian approach used, it is worth highlighting that incorporating enviromic sources does not replace the incorporation of a design matrix for environments (here used as fixed effects) as it is commonly associated in previous studies of GP reaction-norm. Here we show that enviromic sources came up as tentative to capture the envirotype-phenotype covariances. The cross-validation scheme used in *Case 1* allowed us to observe that the joint prediction of different genotype-environment conditions (Fig 3) might better highlight how enviromic sources can contribute to increasing the predictive ability of GP, mostly due to its usefulness in approaching the environmental correlation among field trials. It shows more transparency for the influence of the scenarios G,*n*E and *n*G*n*E, in which we had a considerable lack of phenotypic information in training GP. We can infer that schemes such as CV1 (only *n*G,E) are the least adequate option to show the benefits of coupling enviromics in GBLUP. However, looking at a drastically sparse MET condition (joint prediction scenarios) shows that enviromics improves the accuracy of GP as the size of the MET also increases. Predictions are made up of tiny experimental networks.

### 4.3 Differences in using environmental covariables (W) and typologies (T)

Regarding the enviromic assembly approaches used in this study, there was evidence that using typologies as envirotype descriptors (**T** matrix) is more biologically accurate in representing environmental relatedness than quantitative descriptors (**W** matrix) based on quantile covariables. This increase in biological accuracy was reflected in the statistical accuracy and then boosted plant breeders’ ability to carry out selections across multi-environment conditions. Further efforts in this sense must be devoted to increasing the level of explanation of the genotype-envirotype covariances, which can also take advantage of Shelford’s Law to refine the limits of tolerance for particular genotypes. Thus, different genotypes will be under the influence of a diverse set of envirotypes, which can be realized for the same environmental factor (e.g., solar radiation, air temperature, soil moisture) according to its occurrence across crop lifetime (e.g., vegetative stage) and the adaptation zone designed from ecophysiology concepts (e.g., temperature cardinals defining which temperature level results in stress and optimum growing condition).

A second difference may be explained by the fact that quantitative environmental covariates are not an additive effect to compose an environment variation. Despite this, we agree with Resende et al. (2020), and we adapted the idea of envirotypes as markers of environment relatedness in a different manner. For example, the common use of mean values of covariates such as rainfall, solar radiation, and air temperature, in reality, represents a non-additive between each other; yet, they are very well correlated for a given site-planting date condition, even when using strategies to deal with collinearity, such as partial least squares (e.g., Vargas *et al*., 2006; Porker *et al.,* 2020;). We can use an example as a given day of crop growing in which a large amount of rainfall has occurred. We can suppose that the sky is cloudy, with less radiation and lower temperature. Thus, using such G-BLUP inspired approach is not an ideal solution to estimate the environmental variance. Conversely, the environmental typologies (**T**) are based on frequencies (ranging from 0 to 1), where the sum of all frequencies are equal to 1 (100% of the variation). In addition, those typologies can be built for a given site using historical weather data, adapting the approach of Gillberg et al. (2019) and de los Campos et al. (2020). As presented in section 2.4.2, if no typologies are considered, the expected environment effect is given for a fixed-environment intercept (with 0 variance within and between environments). Despite this fact, another option is using nonlinear kernel methods to estimate only the environment-relatedness, as this approach takes advantage of nonlinear relationships among covariates (Costa-Neto et al., 2021a,b).

### 4.4 Does more phenotype data mean more accuracy in multi-environment prediction?

This study shows that environmental information can break the paradigm that claims that more phenotype information leads to greater accuracy of GP models over MET. Our results highlight that the traditional GBLUP models assume that the variation due to G×E is purely genomic-based across field trials, leading to an implicit conclusion that the yield plasticity is constant (slope ∼ 0) for all genotypes, which is unrealistic. It also reflects that G×E patterns are non-crossover (scale changes in performance across different variations), that is, a well-performing genotype will always be good across environments, and a poorly performing genotype has the same trend for all environments. Despite the gains achieved in predicting the quality of a novel environment and the plasticity for tested and untested genotypes, we noticed that the inclusion of enviromic sources also leads to the unrealistic conclusion that all genotypes respond in the same way the gradient of climate and soil quality. Our results show a reasonable accuracy in predicting yield plasticity, but further efforts must be made to improve this approach’s explanation of the yield plasticity as a nonlinear variation across the gradient of environmental factors.

The use of selective phenotyping strategies made up with enviromic assembly × genomic kinships showed a drastic reduction of in-field efforts. Combined with enviromic-aided GBLUP models, it led to almost the same predictive ability achieved using a wide number of genotypes and environments for a large experimental network. Thus, we can enumerate the benefits of the enviromic approaches tested in this study as (1) the possibility of training prediction models for yield plasticity with reduced phenotyping efforts, (2) a consequence of the assembly of enviromics with genomics allowing the selection of the genotype-environment combinations that best represents the main inner covariances among phenotypes produced by different environments (the genotype-phenotype, envirotype-phenotype dynamics mentioned above).

Considering both enviromics approached, we conclude that the advantages of E-GP over W-GP can be enumerated as (1) the flexibility to design a wide number of environment-types assuming different frequencies of occurrence of key stressful factors in crop development; (2) it allows the use of historical weather and in-field records to compute trends of certain envirotypes at certain environments, which can be coupled into (3) the definition of TPE and characterization of mega-environments, as the main approach used for this relies on the study of the frequency of occurrence of the main environment-types (e.g., Heinemann *et al*., 2019). For the latter, for example, the **T** matrix proposed here is just an arrangement of an environment × typology matrix, in which each entry represents its frequency of occurrence at a particular time interval of the crop lifetime. Conversely, the advantages of W-GP over E-GP rely on plasticity in creating large-scale envirotype descriptors with reasonable biological accuracy.

### 4.5 Can we envisage climate-smart solutions from enviromics with genomics?

Modern plant breeding programs must deliver climate-smart solutions cost-effectively and time-reduced (Crossa et *al.*, 2021). By climate-smart solutions, we mean (1) adopting cost-effective approaches capable of providing fast and cheap solutions to face climate change (2) a better resource allocation for field trial efforts to collect representative phenotype information to feed prediction-based platforms for crop improvement, such as training accurate GP models and CGM-based approaches capable of guiding several breeding decisions, (3) a better understanding of which envirotypes most limit the adaptation of crops across the breeding TPE, revising historical trends and expecting future scenarios (e.g., Ramirez-Villegas *et al*., 2018; 2020; Heinemann *et al*., 2019) (4) understanding the relationship between secondary traits and their importance in explaining the plant-environment dynamics for given germplasm at given TPE (e.g., Cooper *et al*., 2021). However, most of those objectives will be hampered if the MET-GP platforms do not consider models with a higher biological meaning (Hammer *et al*., 2019) and reliable environmental information. A cost-effective solution for that, if the breeder has no access to sensor network tools, relies on the use of remote sensing tools to collect and process basic weather and soil data, such as those available in the *EnvRtype* R package (Costa-Neto *et al*., 2021b).

If selective phenotyping is added in the enviromics-aided pipeline for GP (Supplementary Fig 1), additional traits and the possibility of screening genotypes across a wide number of managed environments will increase. It can support field trials’ training for CGM approaches, which demands phenotyping of traits across crop life, such as biomass accumulation and partitioning among different plant organs. Finally, using models considering an explicit environmental gradient of key-environmental factors is a second alternative for this approach. It can be done to discover the genetic determinants of the interplay between plant plasticity and environment variation. As a wide range of genes reacts to each gradient of environmental factors, the use of whole-genome regressions of reaction-norm for each environmental factor must be useful to screen potential genotypes (in our case, single-crosses) for a diverse set of scenarios (e.g., increased heat stress). Pioneer works used this methodology in wheat breeding (Heslot *et al*., 2014; Ly *et al*., 2018) inspired other cereal crop applications.

For example, Millet *et al*. (2019) fine-tuned the methodology by creating a two-stage analysis of factorial regression (FR) involving environmental data, followed by a GP based on the genotypic-specific sensibility for key environmental factors found in the FR step. In general, studies involving FR analysis found that the effect of high temperatures at grain-filling and maturation (Epinat-Le Signor *et al*., 2001; Romay *et al*., 2010), water balance at flowering (Epinat-Le Signor *et al*., 2001; Millet *et al*., 2019) and intercept radiation at the vegetative phase (Millet *et al*., 2019) are the main drivers of G×E for yield components in maize. Thus, Millet *et al*. (2019) explores this opportunity offered by FR to use genotypic-specific regressions, which coupled with genomic data, led to an increase of the accuracy of MET-GP by 55% concerning the benchmark environmental similarity model made up of mean values of environmental factors, as proposed by Jarquín *et al*. (2014).

From the aspects mentioned above, we envisage that the use of GP for multi-environment predictions must account for some degree of ecophysiological reality while also considering the balance and the relation between parsimony and accuracy (Hammer *et al*., 2019; Costa-Neto *et al*., 2021b; Cooper *et al*., 2021). Here we also highlight in our literature review that multi-environment GP must account for the impact of (1) resource availability in the creation of biologically accurate platforms in training CGM-based approaches and delivering reliable envirotyping information for those purposes, (2) availability of the knowledge of experts in training CGM approaches. Thus, ecophysiology concepts to provide solutions for raw environmental data processing in enviromic assembly information for predictive purposes seem to be a cost-effective alternative to leverage accuracy involving parsimony and biological reality.

## Supporting information

Suplementary Contents

