## Supplementary material for "Enviromic assembly increases accuracy and reduces costs of the genomic prediction for yield plasticity": Suplementary Contents

### 1 Supplementary Tables

**Supplementary Table 1**. Geographic coordinates of each experimental site and environments used in Multi-Regional Set in 2015, South America, Brazil.

| ID | Site | Region | Year | Management | Latitude | Longitude |
|---|---|---|---|---|---|---|
| NM | Nova Mutum | Middle-Western | 2015 | all necessary management practices to achieve Potential Yield | -13.05 | -56.05 |
| SO | Sorriso | | | | -12.32 | -55.42 |
| PM | Patos de Minas | Southwestern | | | -18.34 | -46.31 |
| IP | Ipiaçú | | | | -18.9 | -49.56 |
| SE | Sertanópolis | South | | | -23.03 | -51.02 |

**Supplementary Table 2**. Geographic coordinates of each experimental site and environments used in N-level set between 2016 and 2017 in South America, Brazil.

| **ID** | **Site** | **Region** | **Year** | **Management** | **Latitude** | **Longitude** |
|---|---|---|---|---|---|---|
| 1_PI_LN | Piracicaba | Southwestern | 2016 | Low N Level | -22.705 | -47.637 |
| 1_PI_IN | | | | Ideal N Level | -22.705 | -47.637 |
| 1_AN_LN | Anhumas | | | Low N Level | -22.87 | -47.997 |
| 1_AN_IN | | | | Ideal N Level | -22.87 | -47.997 |
| 2_PI_LN | Piracicaba | | 2017 | Low N Level | -22.705 | -47.637 |
| 2_PI_IN | | | | Ideal N Level | -22.705 | -47.637 |
| 2_AN_LN | Anhumas | | | Low N Level | -22.87 | -47.997 |
| 2_AN_IN | | | | Ideal N Level | -22.87 | -47.997 |

**Supplementary Table 3.** Estimated variance components (±standard deviation) for N-level set.

| Effect | Model | | | | |
|---|---|---|---|---|---|
| | GBLUP | W-GP (BD) | E-GP (BD) | W-GP (RN) | E-GP (RN) |
| T | - | - | 0.195±0.016 | - | 0.324±0.029 |
| W | - | 0.319±0.029 | - | 0.643±0.06 | - |
| A | 1.371±0.031 | 1.392±0.03 | 1.350±0.03 | 1.330±0.029 | 1.396±0.036 |
| D | 0.574±0.005 | 0.575±0.005 | 0.575±0.005 | 0.550±0.005 | 0.426±0.005 |
| AE | 0.363±0.003 | 0.366±0.004 | 0.365±0.003 | - | - |
| DE | 0.337±0.003 | 0.338±0.003 | 0.336±0.003 | - | - |
| AT | - | - | - | - | 0.034±0 |
| AW | - | - | - | 0.016±0.001 | - |
| DT | - | - | - | - | 0.012±0 |
| DW | - | - | - | 0.006±0.002 | - |
| Residual | 1.154±0.004 | 1.153±0.004 | 1.155±0.004 | 1.45±0.003 | 1.37±0.003 |
| Explained variance (%) | 70% | 72% | 71% | 64% | 62% |

**Supplementary Table 4.** Estimated variance components (±standard deviation) for Multi-Regional Set.

| Effect | Model | | | | |
|---|---|---|---|---|---|
| | GBLUP | W-GP (BD) | E-GP (BD) | W-GP (RN) | E-GP (RN) |
| T | - | - | 0.157±0.019 | - | 0.055±0.004 |
| W | - | 0.090±0.007 | - | 0.078±0.007 | - |
| A | 0.464±0.012 | 0.465±0.012 | 0.464±0.011 | 0.462±0.011 | 0.499±0.014 |
| D | 0.218±0.004 | 0.217±0.004 | 0.216±0.004 | 0.211±0.004 | 0.232±0.005 |
| AE | 0.197±0.003 | 0.199±0.003 | 0.195±0.003 | - | - |
| DE | 0.089±0.001 | 0.088±0.001 | 0.089±0.001 | - | - |
| AT | - | - | - | - | 0.007±0.009 |
| AW | - | - | - | 0.004±0.002 | - |
| DT | - | - | - | - | 0.002±0.001 |
| DW | - | - | - | 0.002±0.003 | - |
| Residual | 0.234±0.001 | 0.235±0.001 | 0.235±0.001 | 0.239±0.001 | 0.237±0.001 |
| Explained variance (%) | 81% | 82% | 83% | 76% | 77% |

### 2 Supplementary Figures

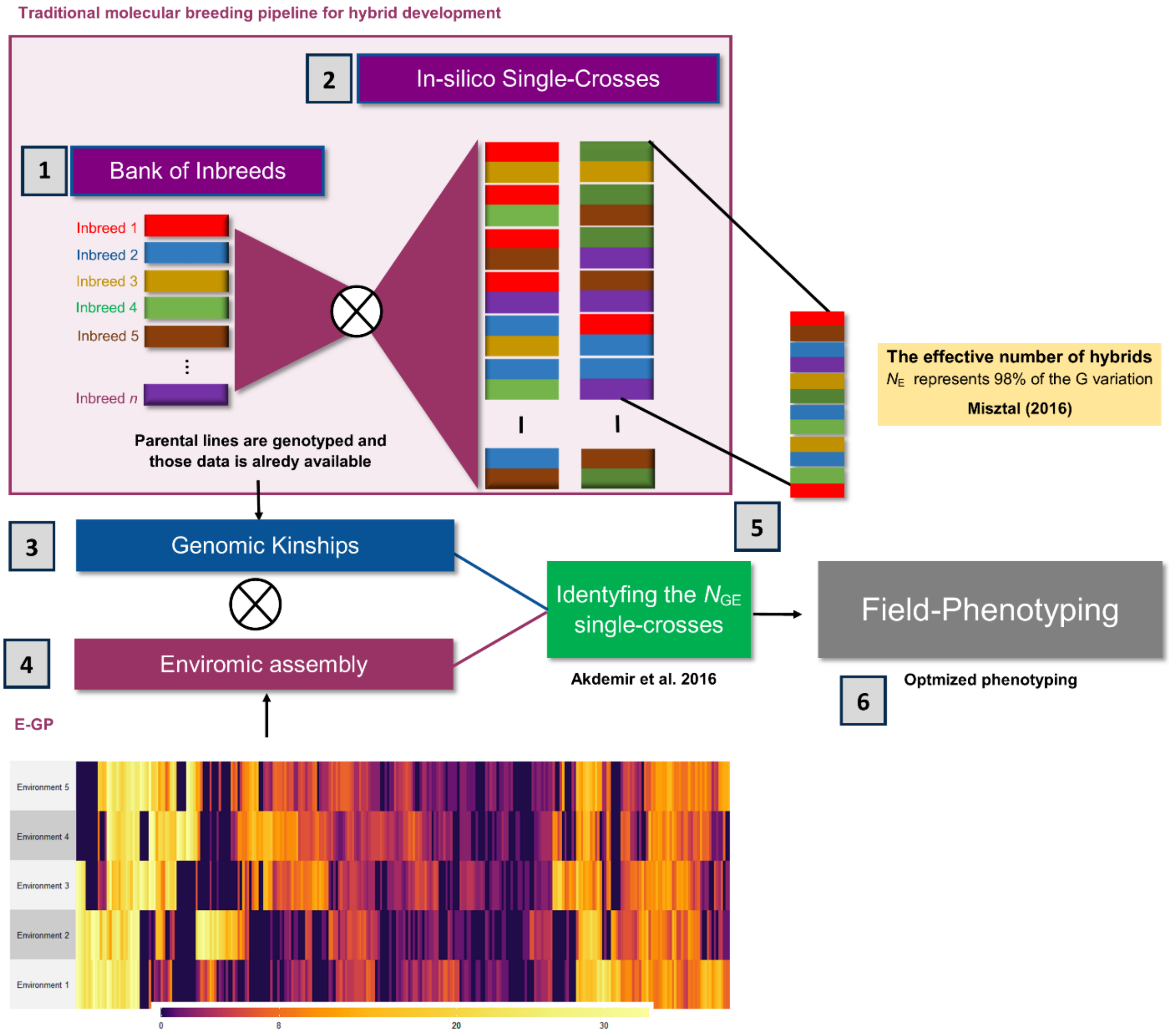


**Supplementary Figure 1. Use of selective phenotyping with genomic kinship and enviromic assembly for boosting the hybrid breeding pipelines.** In purple, it is presented the current molecular breeding approaches of hybrid development. (1-2) **(1)** From a bank of elite inbreeds already genotyped, it is possible to create a large number of in-silico single-crosses **(2)** using the Kronecker product between each SNP marker from desirable parentals. Then, a single-value decomposition (SVD) of the genomic kinship (G) reveals the effective number of genotypes ($N_E$) that represents at least 98% of the variation in G **(3)**. However, under the E-GP platform, the use of G plus an enviromic assembly (T) for a target population of environments (TPE) can be used to build up in silico possible growing conditions that crops may experience **(4)**. Then, via the Kronecker product between enviromic x genomic, it is possible to build a matrix accounting for genotypic observations per environment. Then, we apply SVD on that to select the effective number of genotypes per environments ($N_{GE}$) that represents at least 98% of the variation of the realized experimental network **(5)**. Later, using the SPTGA package, that provides a genetic algorithm, we can define the most relevant combinations of genotype x environment **(6)**. Finally, only these individuals are phenotyped in specific locations. Then, it will be used as a training population for genomic-based prediction or other research purposes, such as training crop growth models or running a factorial regression analysis. This approach allows an optimized training of those models, which may increase efficiency in predicting phenotypic landscapes across novel growing conditions

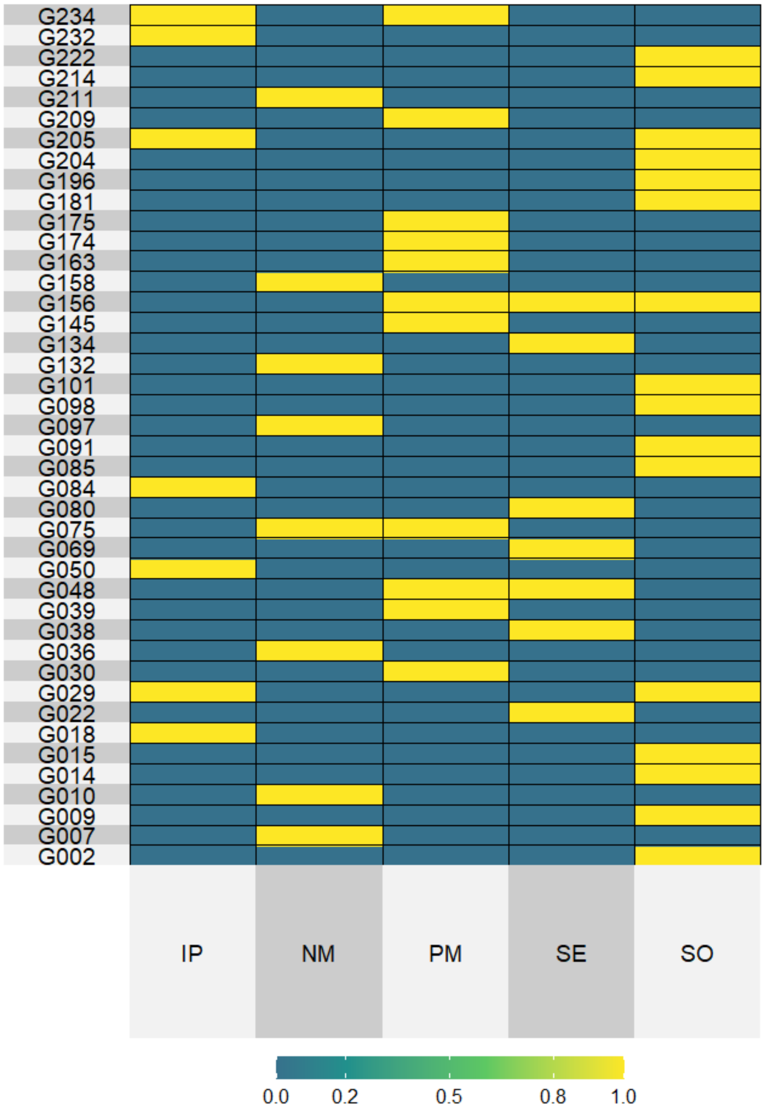


**Supplementary Figure 2. Summary of the selective phenotyping approach drawn by the effective number of observations ($N_{GE}$) phenotyped in the field for the Multi-Regional Set (247 tropical maize hybrids over 5 locations).**The core of 42 maize hybrids (rows) per 5 environments (columns). In yellow is the hybrid-environment combinations phenotyped in the field-based trials. It resulted in $N_{GE}$ = 49, because some genotypes occur in more than one environment. At each environment, the number of genotypes was: IP (7), NM(8), PM (11), SE (7), and SO (16). The remaining 205 hybrids plus the blue cells were considered a testing set (virtual experimental network).

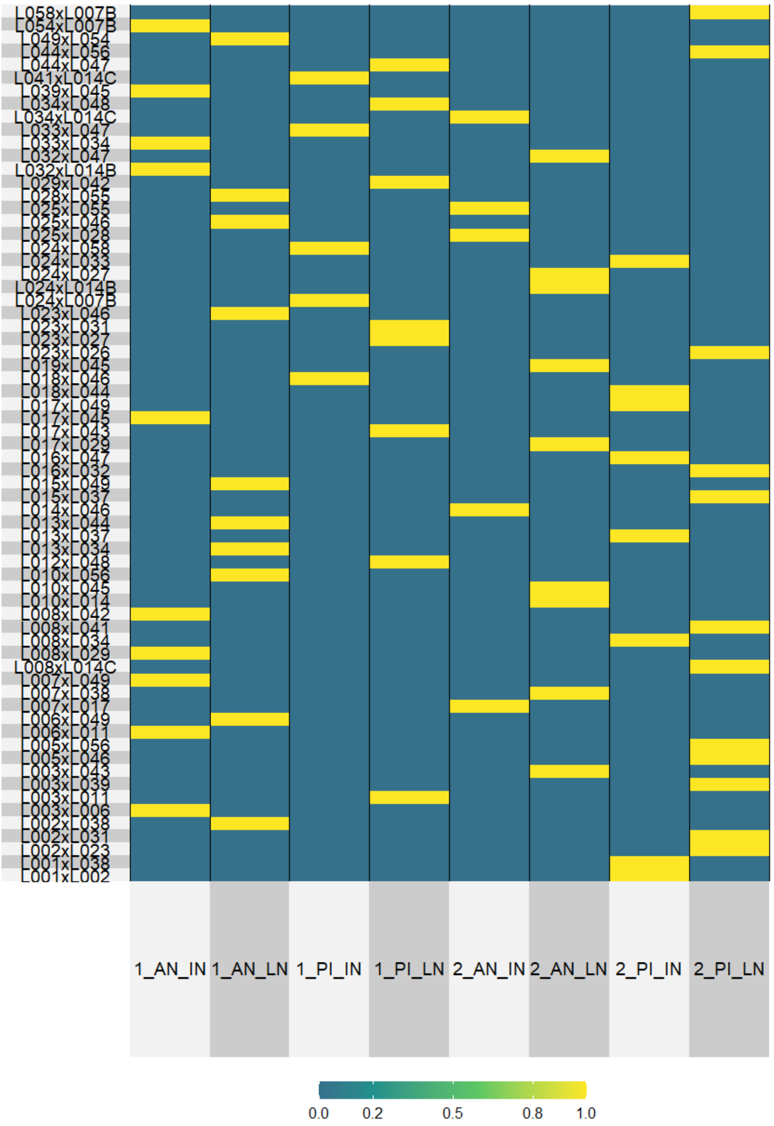


**Supplementary Figure 3. Summary of the selective phenotyping approach drawn by the effective number of observations ($N_{GE}$) phenotyped in the field for the N-level Set (570 tropical maize hybrids over eight environments).** The core of 67 maize hybrids (rows) per 8 environments (columns). In yellow is the hybrid-environment combinations phenotyped in the field-based trials. It resulted in $N_{GE}$ = 67. At each environment, the number of genotypes were: 1_AN_IN (10), 1_AN_LN (10), 1_PI_IN (5), 1_PI_LN (8), 2_AN_IN (5), 2_AN_LN (9), 2_PI_IN (8), 2_PI_LN (12). The remaining 503 hybrids plus the blue cells were considered as testing set (virtual experimental network).
